# I helix Mediates the Allosteric Regulation in Cytochrome P450cam

**DOI:** 10.1101/2025.02.06.636992

**Authors:** Mohammad Sahil, Jagannath Mondal

## Abstract

Cytochrome P450cam, a key monooxygenase in the P450 superfamily, is pivotal in metabolic and industrial processes. Despite extensive studies, a unified mechanism governing its conformational heterogeneity, substrate-dependent allostery, and multi-substrate binding remains elusive. Here, integrating molecular dynamics simulations, NMR pseudocontact shift (PCS) analysis, and crystallographic data, we identify the I-helix (*α*I) as the central regulator of P450cam’s allostery. Its intrinsic flexibility, dictated by glycine residues (G248 and G249), orchestrates enzyme conformational dynamics. Specifically, *α*I transitions between *straight* and *kinked* conformations, modulating the opening and closing of substrate access channels (channel-1 and channel-2) and mediating allosteric communication between active and allosteric sites. Substrate binding stabilizes the *straight* conformation, promoting channel closure and enhancing allosteric regulation. This I-helix-based mechanism reconciles 125 crystallographic poses, spanning straight-to-kinked *α*I conformations. Notably, the kink-inducing glycine G249 is evolutionarily conserved across species, including humans, underscoring *α*I’s fundamental role in enzyme function and broader significance within the P450 superfamily. NMR PCS measurements align with the kinked and straight conformations in the substrate-free and substrate-bound states, with Q-scores of 0.108 and 0.061, respectively. Leveraging this mechanistic insight, we designed proof-of-concept P450cam mutants locked in either constitutively open or closed conformations for the first time. By shifting the focus from the traditional FG-helix-centric view to an I-helix-centric framework, this study provides a comprehensive blueprint for conformational and allosteric regulation, paving the way for engineering tailored P450 variants.

## Introduction

Cytochrome P450 enzymes (CYPs) constitute one of the most versatile families of heme-containing monooxygenases, catalyzing diverse oxidative reactions across metabolic and biosynthetic pathways.^1,2^ Their unparalleled ability to functionalize inert substrates makes them indispensable in pharmaceutical development, industrial biocatalysis, and natural product synthesis.^3,4^ The overall chemistry of heme-reaction center and P450 structural fold are largely conserved in entire P450 superfamily. Among CYPs, cytochrome P450cam (CYP101A1) is a prototypical member, extensively studied for its structural and mechanistic insights into the superfamily.^1^ As the first CYP to be characterized structurally,^5^ P450cam serves as a model system for elucidating enzyme-substrate interactions and understanding the dynamic conformational changes critical for function.

Central to P450cam’s function are its dynamic substrate access channels, channel-1 and channel-2, which regulate the entry of substrates to its heme active site and exit of products (Figure 1). The channel-1 is posited to explore open conformation in absence of substrate^6–8^ thus allowing the latter to enter the active site.^9^ Upon binding to substrate, the channel-1 transition to closed conformation so that the reaction can happen in a sealed environment.^10–12^ Similarly, channel-2 explores open conformations in absence of substrate and closed conformation in presence of substrate. ^13^ Together both the channels exhibit substrate dependent conformations via allosteric controls.^14^

**Figure 1:**
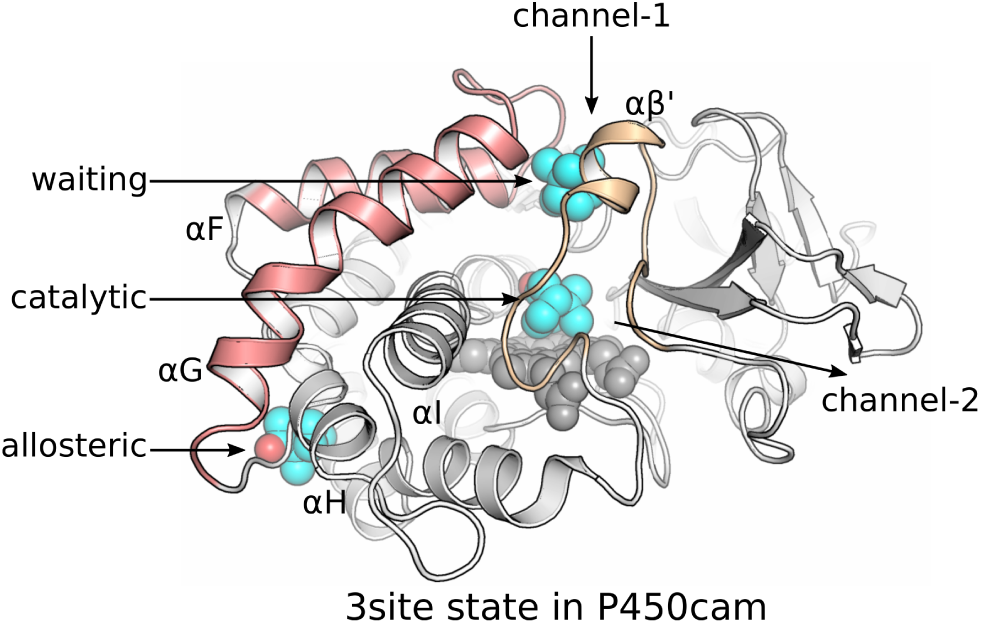
The P450cam. The cytochrome P450cam with highlighted channel-1, channel-2 and its three binding modes. The cyan spheres represents the substrate camphor.

Until recently, only one substrate was known to bind P450cam at its heme active site. However, this enzyme’s lately discovered ability to simultaneously bind copies of substrates at multiple sites both within and outside of heme active site has added layer of complexity to inner working of this protein in the form of allosterically regulated substrate recognition. The single site binding at cataytic heme active site is often contradicted with rare evidences of presence of additional binding modes.^15,16^ Evidence from NMR T1 relaxation times suggested a second binding mode located approximately 15-16 away from the heme Fe atom, involving a conserved threonine residue.^17^ Subsequent molecular dynamics simulations identified a second substrate-binding site at the junction of the E, F, G, and H helices, termed the allosteric site.^18^ This allosteric site was found to be conserved across many P450 enzymes,^19,20^ with crystallographic^21^ and NMR^22^ evidence further supporting its existence. Recently, P450cam was demonstrated to simultaneously bind three substrates: ^13^ the heme active site (“catalytic mode"), the previously proposed allosteric site (“allosteric mode"), and an additional binding site within channel-1 above the heme active site (“waiting mode") (Figure 1). This three-substrate-bound state, referred to as the *3site state*, aligns with NMR pseudocontact shift (PCS) data, suggesting it represents the physiological state of P450cam. The three binding modes within the 3site state adheres to principle of allosteric reciprocity, mutually influencing each other’s thermodynamic and kinetic stabilities. Furthermore, these binding modes function synergistically, with each contributing additively to the functionally critical conformational changes in the active site. ^13^ Similar multi-substrate binding behavior has been observed in other P450 family members, including human variants.^23–27^ This multi-substrate binding not only enhances catalytic efficiency but also introduces a complex layer of allosteric regulation that has been challenging to fully elucidate. While traditional models have emphasized the FG-helix as the primary target of conformational changes, this perspective falls short of explaining the enzyme’s intricate conformational heterogeneity, particularly its allosteric coupling and multi-substrate binding dynamics. Recent discoveries of additional substrate-binding modes and the role of redox partners have only deepened the mystery,^14^ highlighting the need for a unifying framework to understand P450cam’s functional dynamics and especially its allosteric mechanism. The attempts to design P450cam of a desired conformation have not been successful, hence severely limiting its understanding and to utilize its full industrial potential.

Here, we present a transformative perspective, identifying the I-helix (*α*I) as the central allosteric regulator of P450cam. Through an integrative approach combining molecular dynamics simulations, NMR PCS measurements, and crystallographic analysis, we reveal that the *α*I, long overlooked, holds the key to the enzyme’s conformational plasticity. Its intrinsic flexibility, driven by conserved glycine residues, enables it to adopt distinct *straight* to *kinked* conformational extremes that dictate the opening and closing of substrate access channels and mediate allosteric reciprocal communication between the active and allosteric sites. This mechanism not only reconciles decades of crystallography and NMR data but also provides a unified working principle for the enzyme’s substrate-dependent behavior. Moreover, we demonstrate the evolutionary conservation of this mechanism, with the kink-inducing glycine G249 preserved across species, including humans. This conservation underscores the fundamental importance of the *α*I in P450 function and suggests that our findings may extend beyond P450cam to other members of the superfamily. Finally, leveraging this mechanistic insight of allostery, we engineer proof-of-concept P450cam mutants with constitutively open or closed conformations, demonstrating the potential to tailor the enzyme’s dynamics for specific applications.

## Results

### I Helix is Central to Conformational Heterogeneity of P450cam

To investigate the conformational dynamics of P450cam, we initiated our study with a wild-type (WT) substrate-free simulation ensemble, focusing on three well-documented conformational changes (Figure 2A):

1. **channel-1 opening/closing**: mediated by the retraction of the *α*FG away from the heme active site.
2. **channel-2 opening/closing**: driven by the separation of loops *L*83 and *L*102, which flank the *αβ^′^*, and occasionally accompanied by the unfolding of *αβ^′^*.
3. **active-allosteric site coupling**: governed by the shifting of the *α*H, which modulates the volume of the allosteric site, as detailed later.

**Figure 2:**
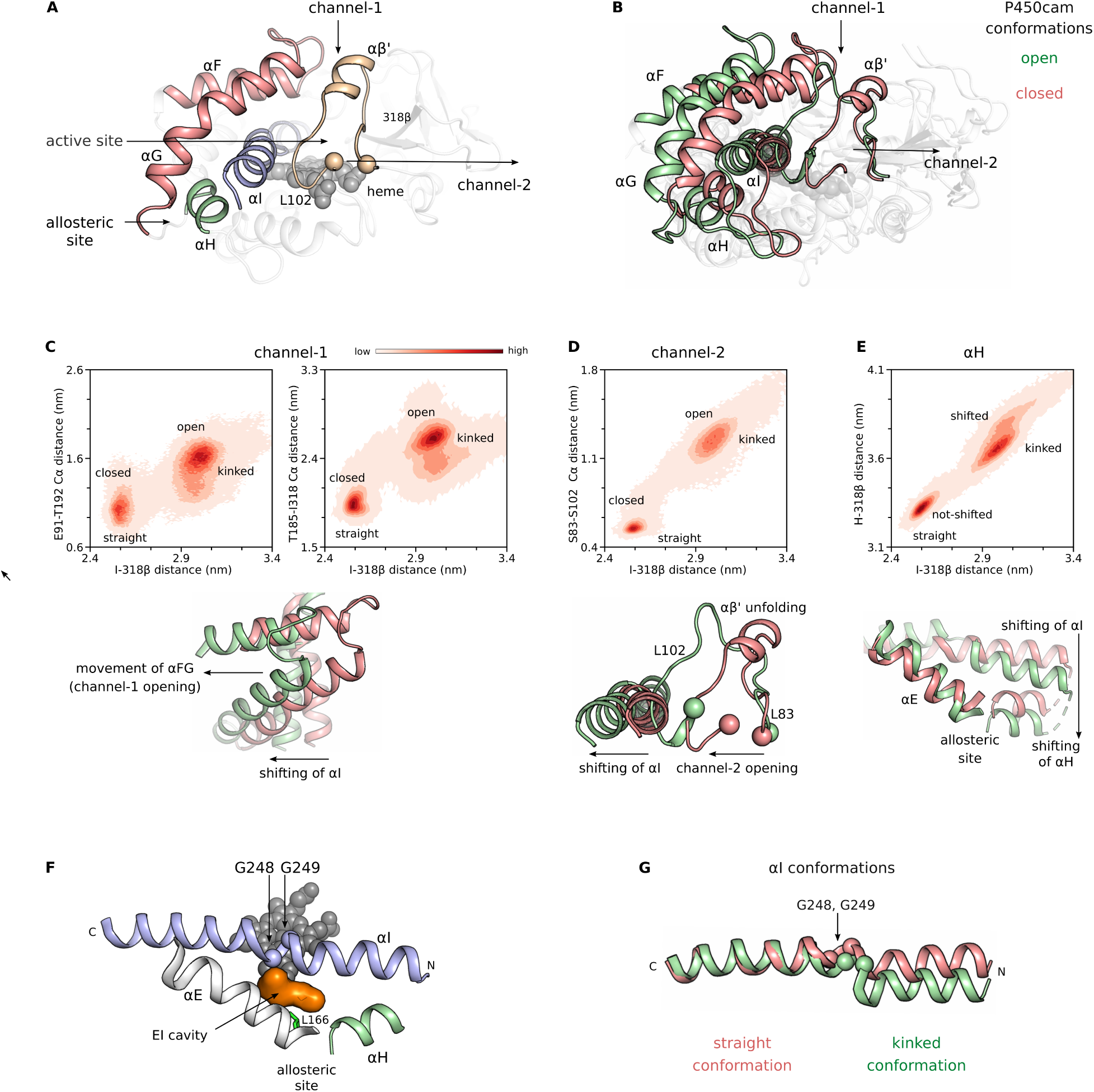
The *α*I as underlying determinant. (A) The cytochrome P450cam in substrate-free state (WT-substrate-free). The main structural elements including channel-1, channel-2, different helices and L102 are color highlighted. (B) The two-ends of conformational spectrum of P450cam observed in WT-substrate-free simulation ensemble. The salmon conformation with straight *α*I, closed channel-1 and channel-2, and non-shifted *α*H. The green conformation with kinked *α*I, open channel-1 and channel-2, unfolded *αβ*’ and shifted *α*H. (C-E) The conformational correlation between conformations of *α*I and channel-1, channel-2 and *α*H. Plots represents the conformational correlation between order parameters of *α*I (x-axis) with that of channel-1, channel-2 and *α*H (y-axis, methods), with red color representing arbitrary probability axis. (F) The zoomed view of *α*I and surrounding environment, highlighting G248, G249 and EI cavity. (G) The zoomed and isolated view of straight and kinked conformation of *α*I.

Analysis of multiple simulation replicates confirmed the characteristic conformational plasticity of P450cam (Figure 2B). Both channel-1 and channel-2 sampled open and closed states, consistent with prior experimental and MD findings. Additionally, the *α*H exhibited a shifted conformation relative to its original not-shifted position (Figure 2B).

A key finding from our simulations was the identification of a fourth, previously unreported, conformational change involving the *α*I (Figure 2B). The *α*I was observed to transition between straight and kinked conformations, with the kinked conformation marked by a deviation away from the heme active site (Figure 2B). Notably, *α*I is strategically positioned within P450cam, structurally linked to all three previously known conformational changes: *α*FG (channel-1), *L*102 (channel-2), and *α*H (allosteric site) (Figure 2). This proximity enabled *α*I to act as a central hub, dynamically coupling these conformational changes. Analysis of the WT-substrate-free simulation ensemble (order parameters in methods) revealed three key conformational correlations:

- the open and closed conformations of channel-1 are directly coupled with the kinked and straight conformations of *α*I respectively. As *α*I shifts to kinked conformation, it pushes *α*FG away from active site, hence opening up the channel-1 (Figure 2C).
- the open and closed conformations of channel-2 are linked to the kinked and straight conformations of *α*I. The kinked *α*I conformation allows *L*102 to separate from *L*83, enabling channel-2 opening (Figure 2D). In contrast, the straight *α*I conformation prevents this separation, maintaining channel-2 in a closed state. While channel-2 can occasionally remain closed despite the kinked *α*I conformation, this scenario is rare in our simulations.
- the shifting of *α*H is tightly coupled with the kinked conformation of *α*I. Due to their sequential and structural proximity, the kinked *α*I induces a corresponding shift in *α*H, with partial involvement of the N-terminal end of *α*E (Figure 2E). These helices are part of the allosteric site and their collective movement bulges the surface of the allosteric site, reducing its depth and volume. For instance, the allosteric pocket volume decreases from 277.45 ± 3.87 Å^3^ (straight *α*I) to 226.59 ± 1.99 Å^3^ (kinked *α*I) (Figure S1).

The *α*I is particularly intriguing due to its intrinsic determinants of conformational heterogeneity, which distinguish it from channel-1, channel-2, and *α*H. First, *α*I contains two glycine residues (G248 and G249) with low helix propensity (Figure 2F). Glycine residues are known to introduce kinks in helices, making *α*I inherently prone to bending. These glycines divide *α*I into two roughly equal segments: residues 234–247 (N-terminal half) and residues 250–266 (C-terminal half). Second, a small cavity (EI cavity) adjacent to the N-terminal half of *α*I provides the necessary space for the helix to shift within the structural fold (Figure 2F).

These intrinsic features enable *α*I to sample straight and kinked conformational extremes, which in turn drive the conformational changes in *α*FG (channel-1), *L*102 (channel-2), and *α*H (allosteric site) (Table 1, Figure 2). Thus, the *α*I helix serves as a central determinant of P450cam’s conformational plasticity, orchestrating the enzyme’s dynamic behavior through its unique structural and conformational properties.

**Table 1:**
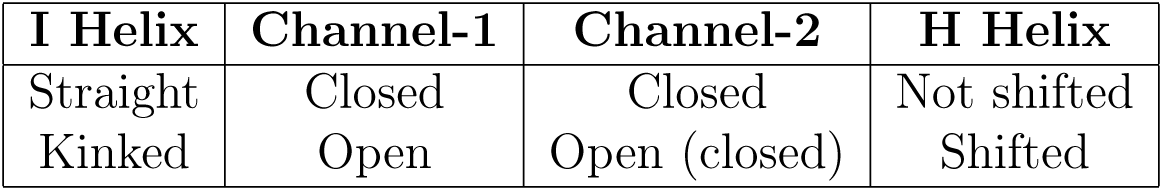
Correlation of *α*I conformations with structural motifs in P450cam. Conformations in parentheses indicate rare occurrences.

### Substrates Control P450cam’s Conformational Heterogeneity via the I Helix

Simulation ensembles of P450cam were generated across all known substrate-binding stages^13^ to investigate their effects on key structural elements (Figure 3). Our analysis revealed that substrates primarily target the *α*I. This helix traverses the active site, where the catalytic binding mode establishes hydrophobic interactions with residues in N-terminal half of *α*I like L244, V247, G248, and G249 (Figure 3A). These interactions partially stabilize the straight conformation of *α*I, thereby limiting its transition to the kinked conformation, as observed in the WT-catalytic simulation ensemble (Figure 3C). On the other hand in allosteric site, the pocket volume is coupled with shifting of *α*H which inturn coupled to transition in *α*I to kinked conformation. The binding of substrate in allosteric site prevents the shifting of *α*H, hence indirectly stabilizing the *α*I in straight conformation (Figure 3A,C). Consequently, the WT-allosteric simulation ensemble displayed a pronounced stabilization of the straight *α*I conformation (Figure 3C). In the presence of the waiting mode-which exists only alongside the catalytic and allosteric modes and stabilizes the catalytic interaction^13^-the hydrophobic interactions between *α*I and the catalytic mode are further strengthened. Together, the three binding modes (the 3site state) synergistically stabilize *α*I in a predominantly straight conformation (Figure 3C). This gradual stabilization aligns with earlier reports that substrate binding modes in P450cam function cooperatively.^13^

**Figure 3:**
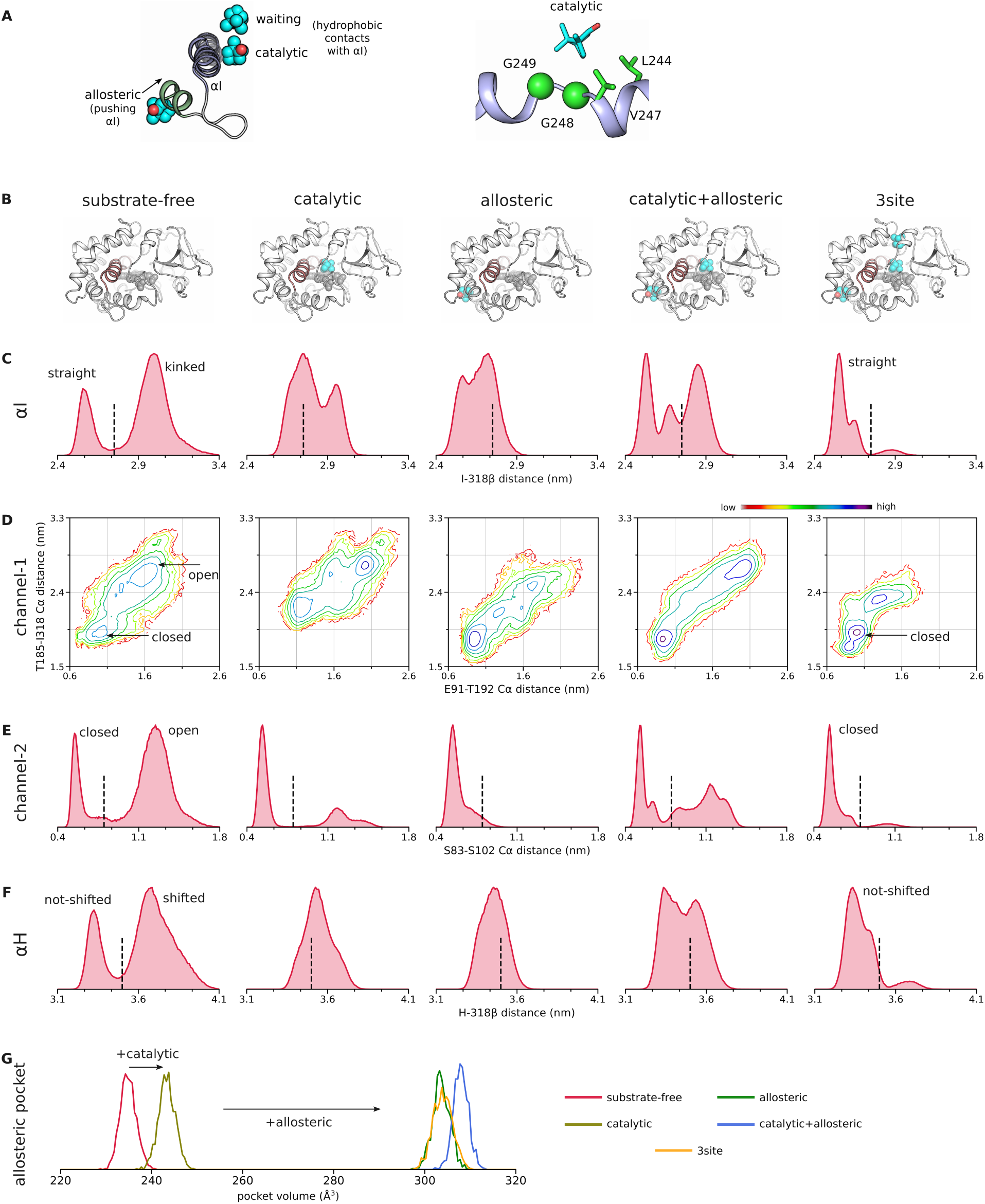
Substrates effector role via *α*I: (A) The interactions of three binding modes of substrates with *α*I. (B) The representative snapshots of progressive substrate binding stages of P450cam. ^13^ The 3site state represents the final bound state. (C-F) The conformations of *α*I, channel-1, channel-2, and *α*H as measured in simulation ensembles of different stages of substrate bindings. The dotted vertical lines in (C,E,F) demarcates their different conformations for comparision. (F) The allosteric pocket volume (^3^) in different stages of substrate bindings. The y-axis in (C,E,F,G) and colorbar in (D) represents arbitrary probability axis.

### Allosteric mechanism

The *α*I possesses intrinsic flexibility, enabling it to transition between straight and kinked conformations. In the substrate-free state, *α*I significantly samples the kinked conformation, which drives the opening of *α*FG and *L*_102_, resulting in open channel-1 and channel-2 conformations. Additionally, this kinked state facilitates the shifting of *α*H. However, as substrates bind and progressively stabilize the straight conformation of *α*I (Figure 3C), channel-1 and channel-2 simultaneously shift toward their closed conformations in direct correlation with *α*I (Figures 3D, E). In the final 3site state, characterized by a predominantly straight *α*I, both channels predominantly exist in closed conformations. This substrate-driven stabilization of *α*I ultimately induces the structural closure of P450cam.

The stabilization of the allosteric binding mode by the catalytic and waiting modes is also mediated by *α*I, which regulates the volume of the allosteric site. Partial stabilization of *α*I in the straight conformation by catalytic binding mode results in a modest increase in allosteric site volume ((substrate-free: 234.84±1.75 ^3^, catalytic:243.35±1.84 ^3^, Figure 3F,G). In the 3site state, the allosteric site is fully expanded (303.97±2.12 ^3^), consistent with prior reports of increased allosteric residence time from 4.1±0.2 *µs* (without active site modes) to 14.1±0.7 *µs* (with active site modes i.e., 3site).^13^

This allosteric expansion reciprocally influences the active site. By restricting the shifting of *α*I, the allosteric mode facilitates the transition of *α*FG toward the closed conformation of channel-1 (Figure 3D). Channel-1 serves as the entry route for catalytic and waiting modes,^9,18^ and its closure stabilizes these modes. This explains earlier findings of enhanced stability in catalytic binding free energy (-5.4±0.3 kcal/mol without allosteric mode vs -7.9±0.6 kcal/mol with allosteric mode) and significantly prolonged residence time of the waiting mode (0.6±0.08 ms without allosteric mode vs 644.9±155.8 ms with allosteric mode).^13^

Overall, the *α*I orchestrates a reciprocal allosteric relationship between the active and allosteric sites. Substrate-induced stabilization of *α*I in the straight conformation regulates allosteric site expansion and closure of channel-1 and channel-2. This, in turn, stabilizes active site binding modes, reinforcing the synergistic coupling of P450cam’s conformational states. Thus, *α*I emerges as a pivotal regulator of the structural and functional plasticity of P450cam.

### Re-examining Available P450cam Data Reconciles the Allosteric Role of the I Helix

In the allosteric mechanism deciphered here, the *α*I emerges as the central and sole allosteric regulator of cytochrome P450cam, an enzyme extensively studied over the past decades. We reasoned that if this mechanism is correct, the straight-to-kinked conformational transitions of *α*I should be observable in crystallographic and molecular dynamics (MD) structures or should leave detectable signatures in NMR spectroscopy and other biochemical or bioinformatics studies. To validate this, we re-examined available NMR, bioinformatics, MD, and crystallography data.

### NMR Evidence

The wild-type (WT) simulations used in this work, representing substrate-free and substrate-bound (3site) states, were previously validated against NMR pseudocontact shift (PCS) data measured in the absence and presence of the substrate camphor.^13^ The simulations matched the NMR data well, as indicated by *C_α_*-*Ln* distances and Q-scores of 0.098 (substrate-free) and 0.045 (substrate-bound), confirming that the *α*I conformations reported here are consistent with NMR PCS measurements. Furthermore, NMR residual dipolar coupling (RDC) studies identified the *α*I, particularly its N-terminal half (residues L244, L245, V247, G248, and G249), as the most perturbed region of P450cam in the presence and absence of camphor.^28–30^ While these observations align with our proposed allosteric mechanism, previous studies attributed these perturbations to *α*I’s role in the active site rather than its allosteric function.

Given that *α*I adopts distinct conformations in substrate-free and substrate-bound states, which drive substrate-dependent conformational changes in P450cam, NMR PCS data measured under these conditions should preferentially match P450cam structures with specific *α*I conformations. According to our mechanism, substrate-free and substrate-bound PCS measurements should align with kinked and straight *α*I conformations, respectively. To test this, we categorized 195001 MD structures from the WT-substrate-free simulation ensemble into 20 groups based on the *α*I-318*β* distance (2.45-3.45 Å), representing the straight-to-kinked conformational spectrum of *α*I. Structures in each category were matched with substrate-free (^15^N-^1^H) and substrate-bound (^2^HN, ^15^N-^1^H) PCS measurements using 5-fold cross-validation (methods). Since not all MD structures are expected to match PCS data due to thermal fluctuations,^22^ we analyzed the fraction of structures that matched the PCS data (benchmarking in Figure S2, Supplementary Note I).

Consistent with our mechanism, substrate-free ^15^N-^1^H PCS data specifically matched structures in the kinked conformational regime, with a mean Q-score of 0.06 (Figure 4A,B). In contrast, substrate-bound ^2^HN and ^15^N-^1^H PCS data matched structures in the straight conformational regime, with mean Q-scores of 0.108 and 0.077, respectively (Figure 4A,B). Similarly, ^15^N-^1^H PCS data for P450cam bound to both substrate and putidaredoxin (pdx) also preferentially matched the straight conformational regime (Figure S3). These results demonstrate that *α*I conformations alone can reconcile substrate-free and substrate-bound NMR PCS measurements, providing direct evidence that substrates modulate P450cam via *α*I.

**Figure 4:**
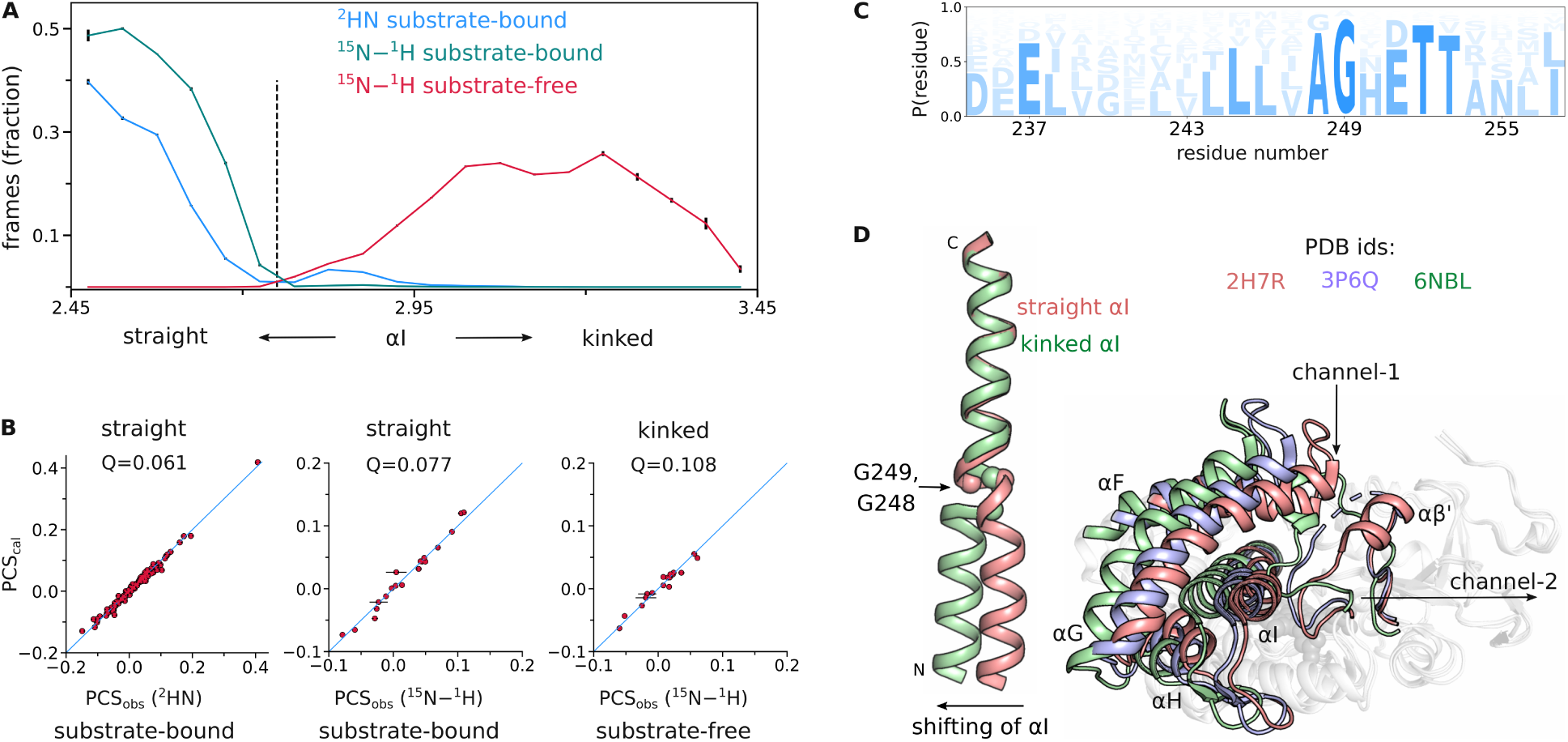
*α*I conformational agreement with NMR and crystals: (A) The fraction of MD structures correctly fitted with different NMR PCS measurements. The vertical dashed line demarcates straight vs kinked conformations of *α*I in accordance with other figures. (B) The matching between observed PCS (x-axis) and calculated PCS from their correctly fitted MD structures (y-axis, top). (C) The concensus sequence of cytochrome P450 family in *α*I region. The residue numbering is according to P450cam (CYP101A1). (D) The crystal structures of P450cam exhibiting straight and kinked conformations of *α*I. The extent of channel-1, channel-2 opening and *α*H shifting are correlated with extent of *α*I kinked conformation.

Revisiting previously reported long MD simulations of P450cam^9,13^ revealed the presence of straight and kinked *α*I conformations and their correlation with channel-1 and channel-2 dynamics. Simulations with closed channel conformations exclusively featured straight *α*I, while those sampling open conformations exhibited kinked *α*I, consistent with our proposed allosteric mechanism (Figure S4).

### Sequence Conservation

To assess the functional importance of G248 and G249, the intrinsic determinants of *α*I conformational heterogeneity, we analyzed their conservation across homologous proteins. Residues critical for function are typically conserved, as seen for the catalytically important D/E251 and T252 (positive controls), which exhibit conservation probabilities of 0.887 and 0.853 respectively, in bacterial P450s. In contrast, G243 (negative control) is not conserved (probability: 0.033). Strikingly, G249 is highly conserved, with a probability of 0.912 across 1,013 sequences^31^ (Figure 4C), supporting its role in P450cam’s conformational heterogeneity. This conservation extends to other kingdoms, including humans, where the corresponding G306 exhibits a conservation probability of 0.819, comparable to D/E308 (0.800) and T309 (0.916) (Figure S5). While G248 is not conserved (probability: 0.158), subsequent analysis (see next section) confirms that G249 alone is sufficient to drive *α*I conformational heterogeneity.

### Crystallographic Evidence

We re-examined 125 available crystal structures of P450cam for evidence of *α*I conformations.^32^ Straight and kinked *α*I conformations were readily identifiable, providing direct experimental support for our findings (Figure 4D, Table S1). For example, structures such as 3L63, 2H7R, and 5WK7 exhibit straight *α*I, while others like 3L62, 4JX1, and 6NBL show varying degrees of kinked *α*I. Importantly, crystals with straight *α*I consistently display closed channel-1 conformations, whereas those with kinked *α*I exhibit open channel-1 conformations (Figure 4D). Similarly, channel-2 opening, *αβ^′^* unfolding, or missing electron density in the *αβ^′^* region are observed only in crystals with kinked *α*I. Additionally, *α*H is shifted in all crystals with kinked *α*I. The extent of channel-1 opening, channel-2 opening, and *α*H shifting correlates directly with the degree of *α*I kinking, as shown in overlays of representative structures (Figure 4D). These observations align perfectly with our proposed allosteric mechanism.

While most crystal structures support our findings, a few (e.g., 1RE9, 3OL5) exhibit either partial (only *α*F) or completely open channel-1 conformations without kinked *α*I, seemingly contradicting our mechanism. However, these structures were artificially opened via linker-bound substrates (Table S1).

Overall, the straight and kinked conformations of *α*I reconcile the entirety of available data and reveal an overlooked aspect of P450cam’s conformational dynamics. With hindsight, previous structural studies targeting P450cam’s conformations have effectively explored the straight vs kinked transitions of *α*I, including NMR (e.g., 3L63 vs. 4JWS,^11^ 2M56 vs. 3W9C^33^), crystallography (e.g., 3L63 vs. 3L62^7^), DEER-crystal (e.g., 4EK1 vs. 3L61^10^), DEER-MD (e.g., 3L63 vs. 4JX1^34^), and MD studies (e.g., 1PHC vs. 3L61,^35^ 4JX1 vs. 2CPP,^36^ 4JX1 vs. 2M56,^37^ 4JX1 vs. 5CP4^38^). Although one NMR study focused exclusively on straight conformations (2L8M vs. 2LQD), ^29^ its findings remains an outlier. While the impact of *α*I has been briefly noted in infrared spectroscopy,^39^ DEER,^34^ NMR,^28^ and crystallography studies, ^21^ this work is the first to assign a functional allosteric role to *α*I.

### Designing Constitutively Closed P450cam with a Straight Variant of the I Helix

We hypothesized that the conformation of channel-1 in P450cam could be stabilized in desired conformation by controlling the conformation of its underlying determinant, the *α*I. To test this, we aimed to design proof-of-concept P450cam mutants that predominantly adopt the straight conformation of *α*I. According to the allosteric mechanism proposed in this study, such mutants are expected to maintain closed conformations of channel-1 and channel-2 even in the absence of substrates, resulting in a constitutively closed P450cam.

As a first approach, we targeted the two key residues, G248 and G249, which are intrinsic sources of the kinked conformation. These glycines were substituted with alanines, which have the highest helix propensities. However, glycines in helices are evolutionarily strategic and challenging to replace. To evaluate the success of these substitutions, we simulated a prototypical subsystem involving only *α*I (residues 234–266) in the wild-type, a constrained system (artificially forced into a straight conformation; see Methods), and various substitution mutants (Figures S6-S7, Supplementary Note II). Mutants that sampled straight conformations similar to the constrained system (positive control) and outperformed the wild-type (negative control) were deemed successful (Figure S6). Trial simulations indicated that the G249W mutant best maintained the straight conformation of *α*I, while G249V also performed reasonably well (Figure S6). The selected mutants were then simulated in the full P450cam system. The G249W substitution is sterically incompatible with the open (kinked) conformation, as confirmed by AlphaFold and RosettaFold predictions, which yielded closed P450cam structures (Figure S7). However, the second-best AlphaFold model resembled the open wild-type structure, so we simulated G249W using both models as starting structures (denoted as G249W-1 and G249W-2).

To assess whether these mutants could achieve substrate-induced closing in the absence of substrates, we simulated them in the substrate-free state. The *α*I in all mutants significantly adopted the straight conformation (Figure 5), in contrast to the binary population of straight and kinked conformations observed in the WT-substrate-free ensemble (Figure 2C). These results demonstrate that the rationally designed mutants effectively reduce the kinking of *α*I to varying degrees. Consistent with the proposed allosteric mechanism, the channel-1 and channel-2 conformations of these mutants sampled closed states corresponding to the degree of *α*I straightening achieved by each mutant (Figure 5C-5D), confirming that channel conformations can be modulated via *α*I.

**Figure 5:**
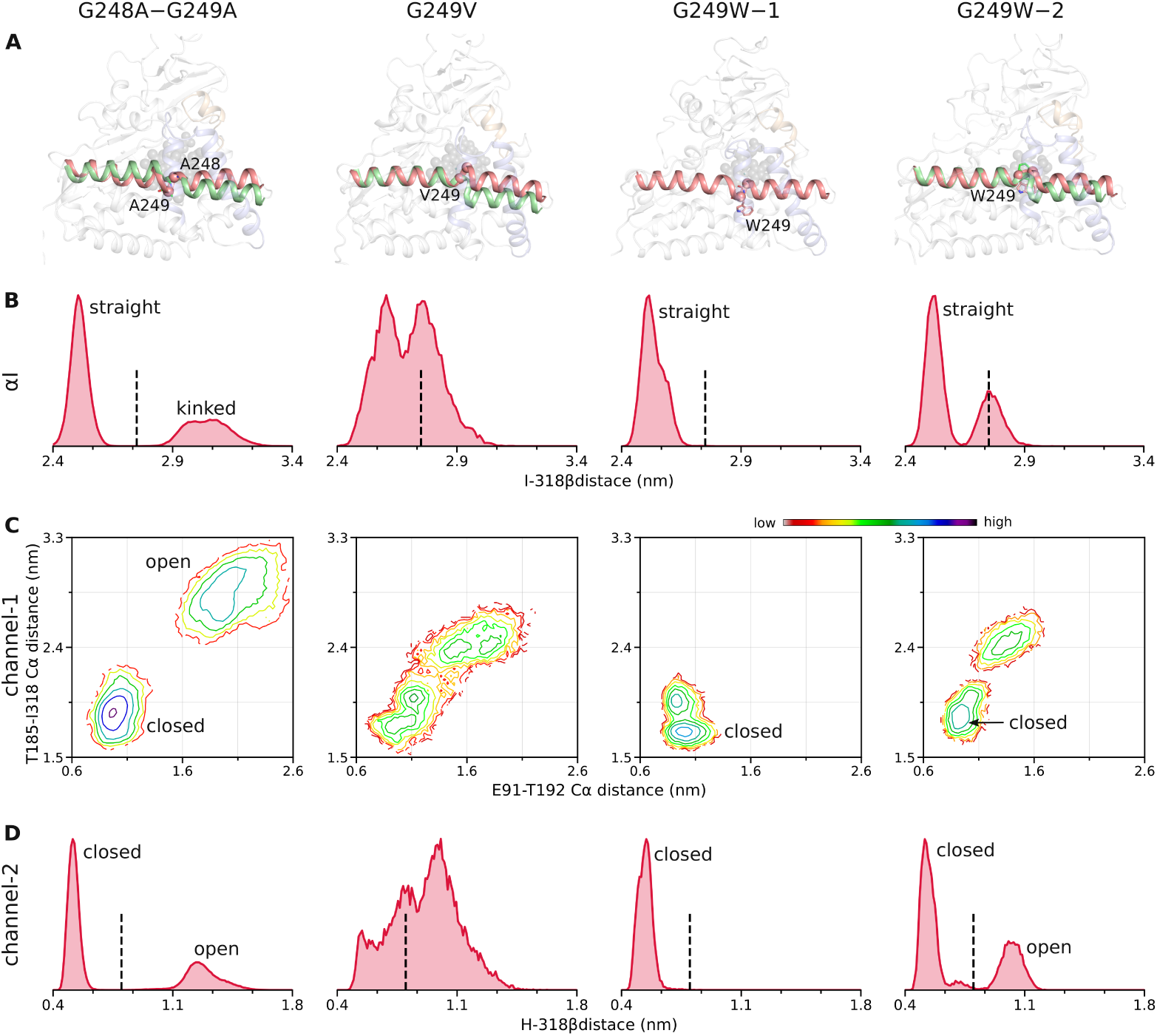
Closed P450cam with straight *α***I**: (A) The P450cam snapshots with *α*I conformations as observed for simulations of different mutants. The snapshots of *α*I are shown for ensembles with multiple conformations. (B-D) The conformations of *α*I, channel-1 and channel-2 as measured in simulation ensembles of different mutants. The y-axis in (B,D) and colorbar in (C) represent an arbitrary probability axis.

In the G248A-G249A substrate-free simulations, *α*I predominantly adopted the straight conformation, with minor kinked conformations also observed. Consequently, the channels primarily sampled closed states, with occasional open states (Figure 5). In the G249V simulations, *α*I sampled straight to intermediately kinked conformations, accompanied by closed-to-intermediate channel conformations. For G249W-1, starting from a closed state, *α*I maintained a predominantly straight conformation, and the channels remained closed. Similarly, in G249W-2, which started from an open state, *α*I significantly shifted toward the straight conformation, resulting in closed channel conformations comparable to those induced by substrate binding in the wild-type simulations (Figure 2). Among the mutants, G249W emerged as the most effective in maintaining a straight *α*I and a constitutively closed P450cam, although its large size may destabilize the heme group in a minor population (Figure S7). As expected, *α*H also sampled the non-shifted conformation to the extent that *α*I was straightened (Figure S8).

### Designing Constitutively Open P450cam with a Kinked Variant of the I Helix

Next, we aimed to design a P450cam variant with a predominantly kinked *α*I. Based on the proposed allosteric mechanism, such a variant would maintain both channel-1 and channel-2 in constitutively open conformations, creating a constitutively open P450cam. Since *α*I already possesses an intrinsic ability to adopt a kinked conformation due to its glycine residues, we sought to stabilize this conformation. We targeted the L166 residue of the *α*E, which forms part of the allosteric site and defines the boundary of the EI cavity that influences *α*I dynamics (Figure 6B-6C). We hypothesized that mutating L166 to a smaller alanine residue would create additional space, favoring the kinked conformation of *α*I. Accordingly, we designed the L166A mutation and subjected it to extensive molecular dynamics simulations.

**Figure 6:**
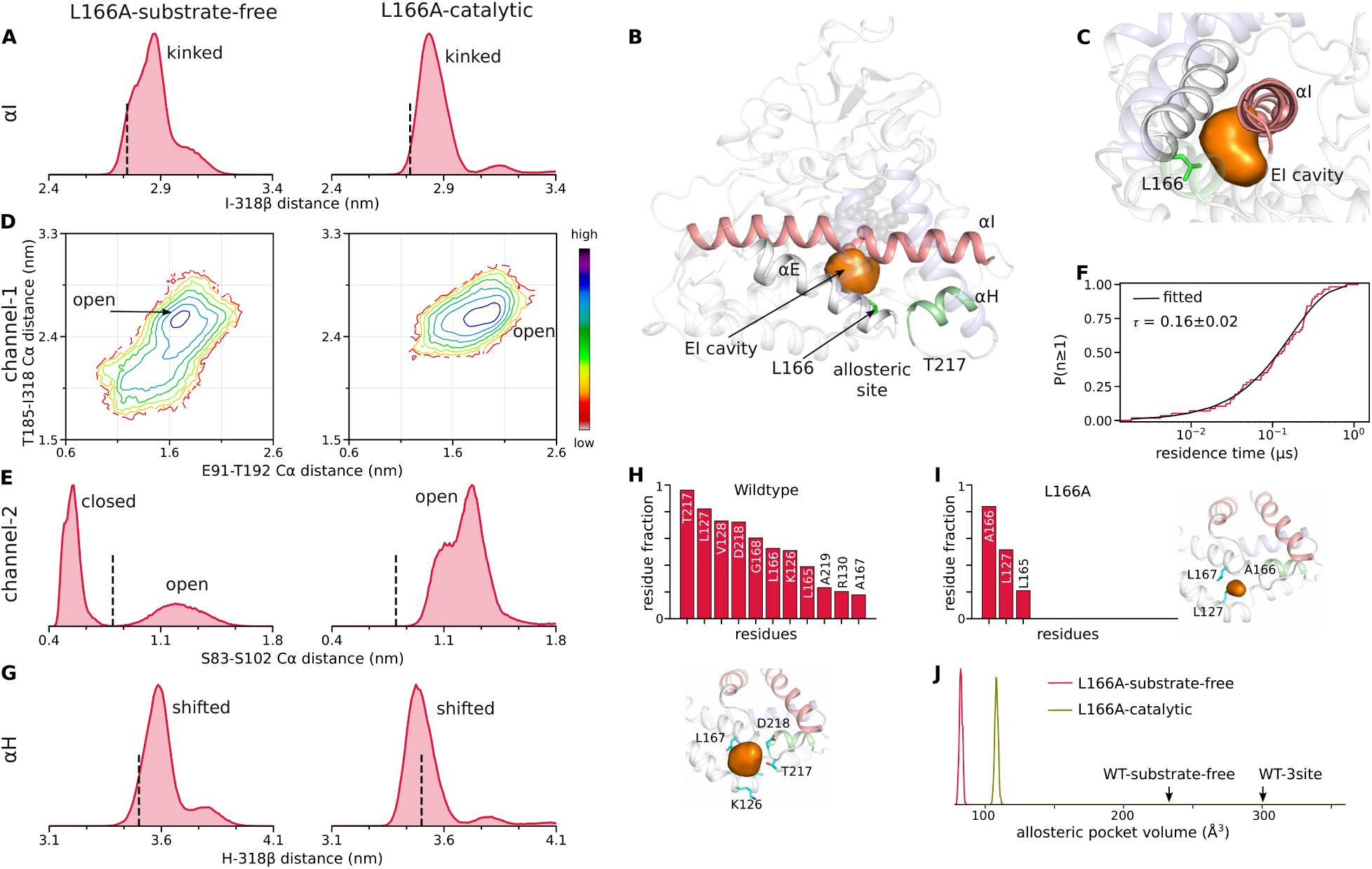
Open P450cam with kinked *α*I: (A,D,E,G) The conformations of *α*I, channel-1, channel-2 and *α*H in L166A mutant P450cam. (B,E) The snapshots of wildtype starting structure highlighting L166 location with respect to *α*I. (F) The infrequent-metadynamics based ECDF of unbinding times. (H,I) The constituent residues of allosteric sites in wildtype and L166A mutant. Snapshots indicate the detected allosteric site in orange surface. (J) The allosteric pocket volume of L166A P450cam. The y-axis in (A,E,G,J) and colorbar in (D) represents arbitrary probability axis.

Consistent with our hypothesis, the L166A-substrate-free simulation ensemble predominantly exhibited the kinked conformation of *α*I, with no occurrence of the straight conformation (Figure 6A). Consequently, channel-1 consistently sampled an open conformation (Figure 6D). Although channel-2 primarily sampled a closed conformation, it occasionally adopted an open state (Figure 6E), aligning with the proposed allosteric mechanism, where channel-2 can remain closed with a kinked *α*I but is sterically capable of opening.

An ideal constitutively open P450cam should resist substrate-induced channel closure. To test this, we investigated whether substrate binding could transition the open L166A-substrate-free state toward a closed conformation. In the L166A-catalytic simulation ensemble, the catalytic substrate failed to stabilize the straight conformation of *α*I (Figure 6A), and the channels remained open (Figure 6D-E). We further examined the allosteric binding mode, which in the wild-type enzyme significantly resists the kinked conformation of *α*I. Intriguingly, the L166A-allosteric and L166A-catalytic+allosteric simulation ensembles could not be generated, as the allosteric site was unable to stably bind the substrate. The allosteric mode spontaneously unbound within hundreds of nanoseconds in all 57 replicates (Figure 6F), with an estimated residence time of 0.16 ± 0.02 *µ*s, representing an 85-fold decrease compared to the wild type (14.1 ± 0.7 *µ*s).^13^ This behavior can be attributed to the significantly reduced size of the allosteric site in the L166A mutant ( 100 Å^3^), which is composed of only three residues (Figure 6H-J).

In summary, the L166A mutant lacks a functional allosteric site and cannot achieve the fully bound 3site state observed in the wild-type P450cam. These findings validate that the proof-of-concept L166A mutant is constitutively open and resistant to substrate-induced channel closure, as substrates can bind only at the heme-active site without inducing conformational changes.

We also explored the design of a decoupled P450cam variant through the F163A mutation, aiming to render channel-1 conformationally independent of *α*I. This decoupling was successfully achieved in the substrate-free state but not in the substrate-bound states (Supplementary Note III, Figure S9). Additionally, we targeted the T217 residue, which forms a hydrogen bond with the allosteric binding mode, by mutating it to valine (T217V). This mutation was intended to disrupt allosteric binding without affecting *α*I dynamics. However, T217V maintained a stable allosteric binding mode with a residence time of 16.5 ± 4.7 *µ*s (Figure S10), supporting the hypothesis that the hydrophobic nature of the allosteric site is the primary determinant of binding stability.

## Discussion

This study uncovers a novel allosteric mechanism in cytochrome P450cam, positioning the *α*I as the central determinant of its conformational plasticity and substrate-induced dynamics. Our findings reveal that the intrinsic ability of the *α*I to transition between straight and kinked conformations, mediated by its conserved glycine residues G248 and G249, underpins the enzyme’s structural and functional heterogeneity. These transitions dynamically couple the *α*I to the active site, channel-1, channel-2, and the allosteric site, providing a unified justification for the long-standing questions surrounding P450cam’s substrate-dependent conformational changes and multi-substrate allostericity.

The *α*I emerges as a key mediator of allosteric reciprocity in P450cam, driving the opening and closing of channel-1 and channel-2 while coupling the active and allosteric sites. Substrates primarily act on the *α*I to stabilize its straight conformation, thereby favoring closed channel conformations and a sealed active site, which are essential for functional efficiency. This mechanism reconciles prior structural and spectroscopic observations, including substrate-free and substrate-bound NMR PCS data^11,33^ and crystallographic^32^ evidence of open and closed channels correlating with kinked and straight *α*I conformations, respectively. Furthermore, the mechanism aligns with multi-substrate binding reports,^13^ wherein catalytic, allosteric, and waiting modes synergistically stabilize the *α*I in its straight conformation, enhancing substrate-dependent channel closure.

The implications of this mechanism extend beyond explaining P450cam dynamics to guiding enzyme engineering. We leveraged these insights to design proof-of-concept P450cam mutants with constitutively closed and open conformations. The G249W and G249V mutants stabilize the straight *α*I conformation, locking channel-1 and channel-2 in closed states and offering models for studying allosteric site interactions without active site interference. Conversely, the L166A mutant promotes a kinked *α*I conformation, keeping channel-1 open and resisting substrate-induced closure, making it a potential model for exploring active site binding independently of allosteric effects. It is noteworthy, that L166A mutant has been studied previously^29^ as long distance effect of substrate, however not via its role on *α*I and allosteric site. These mutants not only validate the proposed mechanism but also demonstrate the feasibility of rationally designing P450cam variants for specific industrial and biochemical applications.

The cytochrome P450 superfamily is distinguished by its conserved structural fold, which spans bacterial to human P450 enzymes. Notably, the location of the heme active site is highly conserved across all P450s, underscoring its fundamental role in catalytic function. The allosteric binding mode initially identified in P450cam has been observed in several other CYPs, including CYP3A4, CYP1A2, CYP2A6, CYP2B6, CYP2C8, CYP2C9, CYP2C19, CYP2D6, and CYP2E1.^19,20^ Similarly, the waiting or waiting-like modes have been identified in various CYPs, such as CYP107 (PDB IDs: 1EGY, 1EUP), CYP158A2 (2D0E), CYP3A4 (2V0M), CYP158A1 (2NZ5), and CYP2C8 (2NNH). Although P450 crystal structures, including those of P450cam, are mostly captured in closed states, the presence of channel-1 or similar openings has been recognized as essential for accommodating large substrates. Such openings have been documented in CYPs like CYP158A1 (PDB ID: 2NZ5). To our knowledge, channel-2 dynamics remain unexplored in other CYP members, highlighting an area for further investigation. Moreover, a kink or bulge in the *α*I, often associated with conserved glycine residues, is evident in nearly all CYP members. This feature is particularly notable in crystal structures such as 2NZ5, 8GK3, and 1W0F. Given that these conformational aspects are widely conserved, it is plausible that the allosteric mechanism described here for P450cam could also govern the dynamics of most, if not all, CYP members. This hypothesis provides a foundation for extending the insights gained from P450cam to the broader P450 superfamily.

From an industrial perspective, the insights provided here have significant implications for the application of P450cam and other CYP enzymes in biocatalysis. P450cam has been utilized for various industrial processes, including economically important reactions like ethane to ethanol conversion,^40^ dehalogenation of endosulfan pollutants, ^41^ directed evolution approaches,^42^ RhFRed biocatalytist^43^ and others.^44^ However, these enzymes often require modifications to meet specific conditions. Understanding the *α*I-driven allosteric mechanism opens new avenues for directed evolution and rational design strategies, enabling the development of tailored CYP variants with enhanced efficiency and substrate specificity. For example, G249W could be employed for selective allosteric-driven processes, while L166A may facilitate reactions requiring open access to the active site. Additionally, these findings could inform future innovations in methane oxidation^45^ and other challenging chemical transformations.^46,47^

Despite the computational nature of this work, the robustness of our conclusions is supported by several factors: triplicate simulations extending to one microsecond, alignment of simulation data with NMR PCS and crystallographic observations, and the ability to reproduce experimentally observed substrate-induced channel closure. While molecular dynamics inherently faces sampling limitations, the agreement between simulation-derived ensembles and experimental data strongly supports the validity of the proposed mechanism. Nevertheless, future studies could complement this work through experimental validation of the proposed mutants.

In conclusion, this study establishes the *α*I as the central regulator of P450cam’s conformational dynamics and its allosteric mechanism. Over the past decades, these questions specifically channel-1 have been the target of numerous studies involving linkered substrates,^48^ DEER,^34^ MD,^18,35^ crystallography,^21^ graph network,^36^ enhanced samplings,^49^ NMR,^28^ mutagenesis,^29^ perturbation response scanning, ^50^ infrared spectroscopy,^39^ their combinations etc and multiple allosteric mechanism/residues have been proposed. However, for the first time to the best of our knowledge, by reconciling decades of experimental and computational data, the current investigation provides a unifying framework for understanding P450cam’s functional plasticity. These insights not only advance the fundamental knowledge of cytochrome P450 enzymes but also pave the way for innovative approaches to enzyme engineering and industrial applications. The *α*I-centric perspective of P450cam offers a transformative lens through which to study and exploit the cytochrome P450 superfamily.

## Methods

### Unbiased MD

Cytochrome P450cam (CYP101A1) was simulated in 19 distinct states, (i) substrate-free: holo protein only; (ii) catalytic: only one substrate in heme active site; (iii) allosteric: only one substrate in allosteric site; (iv) catalytic+allosteric: two substrates in catalytic and allosteric sites; (v) 3site: three substrates in catalytic, waiting and allosteric sites; (vi-ix) L166A in substrate-free, catalytic, allosteric and catalytic+allosteric states; (x) T217V-allosteric; (xi-xiv) G248A-G249A, G249V and G249W (G249W-1: closed, G249W-2: open) in substrate-free states; and (xv-xix) F163A in substrate-free, catalytic, allosteric, catalitc+allosteric and 3site states. Waiting mode was only stable in 3site state,^13^ hence no simulations were performed for waiting, catalytic+waiting and allosteric+waiting states.

For starting structure of all the states, the crystallographic structure (pdb-4JX1,^51^ chain A) was used as template on which, (a) binding modes were introduced based on knowledge from unbiased binding simulations; ^13^ and (b) mutations were modelled using charmmgui,^52^ rosettafold^53^ or alphafold. ^54^ Protonation states of residues corresponds to neutral pH, except D297 in protonated form to account for its hydrogen bond with one of heme propionate moieties. For simulations, the starting structure in desired state was taken in a cubic water box with charged neutralized by *Na*^+^ ions giving a neutral simluation box of side 9.1 nm with around 74k atoms. Protein, heme, ions were parameterized with charmm36 forcefield ^55^ (with backbone CMAP correction), water was charmm-TIP3P^56^ and substrate camphor via GAAMP generated parameters.^9^ Temperature and pressure were maintained at 303.15 K and 1 atm via Nose-Hoover^57,58^thermostat (relaxation time=1 ps) and Parrinello-Rahman^59^ barostat (*τ* =5) respectively. Non-bonded interactions were calculated within 1.2 nm cutoff via PME summation^60^ and verlet^61^ schemes and hydrogen bonds were constrained using lincs^62^ and settle^63^ algorithms.

Simulations were performed in gromacs-20xx package. ^64^ Followed by position restrained energy minimization, NVT (5 ns) and NPT (5 ns) equilibrations; production runs were performed for a maximum of 1 *µ*s time, unless a desired binding mode unbinds. Atleast three independent simulation replicates were performed. Simulations only after 350 ns time were considered for further analysis, hence the simulation ensembles were independent of starting structure. States involving T217V mutants were only ran for 400 ns in twice replicates and only used for generating starting structures for subsequent infrequent metadynamics. States involving L166A mutation and allosteric mode were stopped prematurely as allosteric mode unbinds (results) though repeated for 57 replicates.

Separately, simulations of only *α*I i.e., residues T234-K266 were also performed in 15 different states, (i) wildtype, (ii) G248A-G249A, (iii) G248A-G249P, (iv) G248P-G249A, (v) G248P-G249P, (vi) V247A-G248A-G249A-L250A, (vii) V247A-G248A-G249A, (viii) G249V, (ix) G249W, (x) G249L, (xi) G249I, (xii) G248A-G249V, (xiii) G248A-G249L, (xiv) G248A-G249I, and (xv) contrained (forced backbone hydrogen bonds between L246-L250 and V247-D251 residue pairs). Starting structure was taken in a cubic simulation box with same parameterization as defined above. Three independent simulation replicates were performed for 20 ns each. Simulations from previous studies^9,13^ were also used in this work.

### NMR pseudo contact shifts

Four TROSY-HSQC based solution NMR PCS datasets were used as follows, (i) ^2^*HN* amide PCS of 117 residues measured in presence of substrate camphor, (ii) ^15^*N* −^1^ *H* amide PCS of 17 leucine residues measured for substrate-free P450cam, (iii) ^15^*N* −^1^ *H* amide PCS of 19 leucine residues measured in presence of substrate camphor, and (iv) ^15^*N* −^1^ *H* amide PCS of 19 leucine residues measured in presence of both substrate camphor and redox partner pdx. These residues sparsely cover the entire enzyme structure and the technical details were provided elsewhere by Ubbink and co-workers.^11,33^ Briefly, amide chemical shifts were measured for E195C/A199C/C334A mutant of P450cam labelled with either paramagnetic (Yb) or diamagnetic (Lu) tag through CLaNP-7 attached to C195 and C199 of *α*G. The PCS represents the difference between chemical shifts of paramagnetically and diamagnetically tagged samples.

For MD and crystallography derived structures, the Δ*χ* tensor and position of lanthonide (Ln) ion were determined using fitting process implemented in paramagpy.^65^ Using *C_α_* coordinates of E195 as initial guess, the singular value decomposition grid search was performed within sphere of 10 radius with 10 fitting points per radius. The fitting was further refined through non-linear regression gradient descent, with ultimate objective to minimize the difference between observed and calculated PCS. The fitted position of Ln ion relative to its attached residues (E195, A199) was used as goodness of fit, which should fall within either 7-9 (strict cutoff) or 5.5-10.5 (tolerable cutoff) distance due to steric reasons^66^ (bench-marking in Supplementary note I). The benchmarking was performed on crystal structures corresponding to pdb ids 3L62, 3L63 and 4JWS as used originally,^11^ randomly created atomic distortions and based on previous works.^13,22^ The 195001 MD structures corresponding to WT-substrate-free simulation ensemble were categorized into 20 categories as per order parameter of *α*I (2.45, 2.50, 2.55, .., 3.45), representing straight to kinked conformational spectrum. For each category, PCS fitting was performed for randomly selected 70% struc-tures in a 5-fold cross validation with deviation treated as error. The difference between calculated and observed PCS was measured by Q-score as:

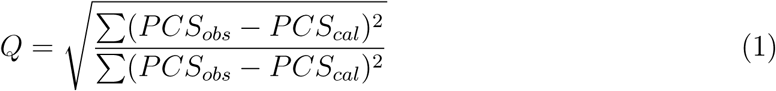

### Crystallographic structures

P450cam crystal structures has been determined with or without substrates and/or redox partner pdx, or with varying factors like -CN bound, oxidation state, *K*^+^ ions, mutations, linkered substrates etc. As or year 2024, total 125 crystal and NMR structures have been determined as per uniprot (P00183) ^32^ (Table S1), which are re-examined in this work.

### Residence time

The residence times for unbinding were estimated for the allosteric mode. For L166A, the allosteric mode unbinds spontaneously repeated for 57 unbiased replicates (results). For other mutants, infrequent metadynamics^67^ approach was used to biasedly unbind the allosteric mode, wereby history dependent gaussian bias potential (*V*_(_*_s,t_*_)_) of height 0.4 kJ/mol and width 0.035 nm was added as a function of distance between substrate and L166. The periodicity of bias was 5000 steps (10 ps) with bias factor of 4, from which unbiased times were re-calculated as eq2 and eq4. The metadynamics was repeated for 50 independent replicates. Unbinding is defined as minimum distance of substrate from 166th residue greater than 1 nm.

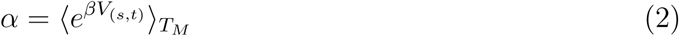

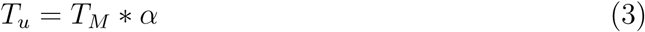

where,

*V*_(_*_s,t_*_)_ is history dependent bias potential as a function of collective variable (s) and time (t). 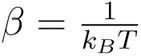 is inverse temperature (T), *k_B_* is Boltzman constant.

Ultimately, a set of *n* unbiased unbinding times 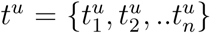 were used to estimate the empirical cumulative distribution function (ECDF, eq4) and was fitted to cumulative distribution function of poisson distribution (CDF, eq5), as unbinding statistics are supposed to follow poisson distribution.^67^ The fitted *τ* represents the residence time. The goodness of fit was confirmed using two sample Kolmogorov-Smirnov test^68^ and p-value greater than 0.05 was considered good (L166A-0.999, F163A-0.179, T217V-0.549). The error was estimated via 100 bootstrapped iterations by taking 70% of observations (with replacement) to fit eq5.

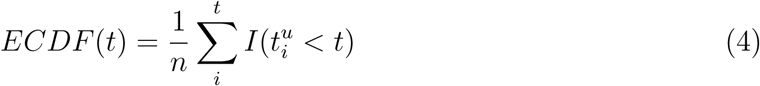

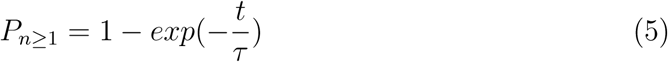

where;

I(.) is indicator function which equals 1 and 0 for true and false argument respectively *P_n≥_*_1_ is probability of observing atleast one unbinding within time t

### Pocket volume estimation

3D pocket volume and constituting residues were estimated for allosteric site on aligned structures taken after every 1 ns from the simulation ensembles. The MDpocket^69^ utility was used to sample the pocket architecture using Voronoi tessellation algorithm. The pocket volume was calculated as cumulative volume of alpha spheres detected in a particular site (allosteric site) for more than 50% of simulation ensemble. In some frames the pocket cannot be detected either due to sidechain flexibility or limitation of MDpocket, such frames were not considered. The pocket volumes of detected frames was bootstrapped for 1000 iterations by randomly iterating with 70% of observations for smoothing of probability density curve and error estimation. The fraction of *i^th^* amino acid (*M_i_*) constituting the allosteric pocket was calculated as weighted sum of its atoms in contact with alpha spheres and normalized by total weighted sum and number of frames as per eq6. The atoms were weighted into different categories i.e., polar (0.5), apolar (0.35) and hydrogens (0.15) or polar (0.59) and apolor (0.41).

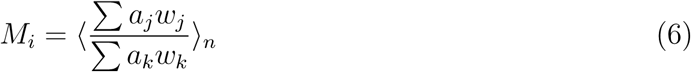

where *a* and *w* are atom types and their weights, *n* is simulation frames, and *j* and *k* represents iteration over only contacted (alpha-spheres) or all atoms of an amino acid respectively.

### Consensus sequence

The bacterial and human P450 sequences were extracted from cytochrome P450 homepage database,^31^ constituting 1151 and 268 sequences respectively. The sequences without gene-id, having large deletions, redundant replicates or not of desired length (300-500 for bacterial, and 400-600 for humans) were not considered, ultimately yielding 1013 and 97 sequences respectively. The alignment was performed using clustalW.^70^ The sequence specific residue probabilities were estimated considering CYP101A1 and CYP3A4 as main sequences, without considering alignment gaps and plotted via logomaker. ^71^

### Order parameters

Channel-1 opening involves motions in *α*F and *α*G, which can sample correlated yet independent conformations. ^13,34^ Unfolding of *αβ^′^* also contributes to channel-1 opening. Hence, channel-1 opening was measured by two independent distances capable of detecting all these motions, i.e., *C_α_* distances between E91(*αβ^′^*)-T192(*α*G) and T185(*α*F)-I318(stable reference) residue pairs. Channel-2 opening/closing is governed by loops preceeding and succeeding *β^′^* helix constituting S83 (*L*83) and S102 (*L*102), whose *C_α_* distances were used as a measure of channel-2 opening, in accordance with previous works.^13,18^ For kink in *α*I, distance between its first half (residue 234-250) and stable reference 318*β* (residues 316-321) was measured as mean of 810 distances corresponding to each possible pair of mainchain atoms. The shifting of *α*H (residue 218-225) was also measured as mean of 432 pairwise distances from stable 318*β*.

## Supporting information

supplemental methods, results and tables

## Supplementary Information

Supplementary figures of pocket volumes, PCS benchmarking, calculated PCS for substrate-bound (+pdx), conformational correlation plots of previous MD data, multiple sequence alignment and consensus sequence, *α*I bending angle, G249W conformations, *α*H in closed variants, F163A decoupled plots and T217V residence time.

Supplementary notes on PCS benchmarking, searching *α*I mutants, and decoupling channel-1 from *α*I.

Supplementary Table of pdb ids of CYP101A1.

## Code and Data Availability

The raw MD trajectories, codes, pocket volumes, sequence aligned outputs, calculated PCS,

residence times and MD parameters are available at zenodo and https://www.github.com/msahilgit/cytoch P450 .

## Acknowledgements

This work was supported by shared computing resources obtained from TIFR Hyderabad, India. We acknowledge support of the Department of Atomic Energy, Government of India, under Project Identification no. RTI 4007. JM acknowledges Core Research grants provided by the Department of Science and Technology (DST) of India (CRG/2023/001426). We thank Dr. Marcellus Ubbink for generously sharing their useful pseudo contact shift data obtained from their NMR measurements on P450cam. We thank Dr. Dheeraj Sarkar for useful discussions.

