## supplemental methods, results and tables for "I helix Mediates the Allosteric Regulation in Cytochrome P450cam"

### Supplementary Information for
### I helix Mediates the Allosteric Regulation in Cytochrome P450cam

Mohammad Sahil* and Jagannath Mondal*

*Tata Institute of Fundamental Research Hyderabad, 36/P Gopanapalli Village, TS-500046, India*

**Contents**

Supplementary Figures S1-10 2-19

Supplementary Note-I 4

Supplementary Note-II 12

Supplementary Note-III 17

Table S1 11

References 20

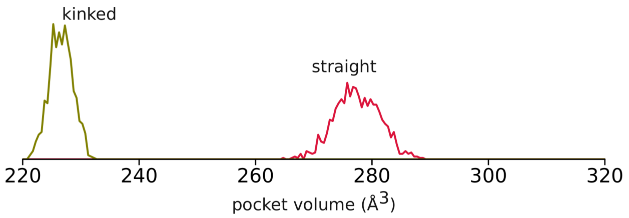


**Figure S1:** The allosteric pocket volume in WT-substrate-free simulation ensemble, measured separately for frames belonging to straight or kinked conformation (demarcated by 2.75 nm I-318$\beta$ distance).

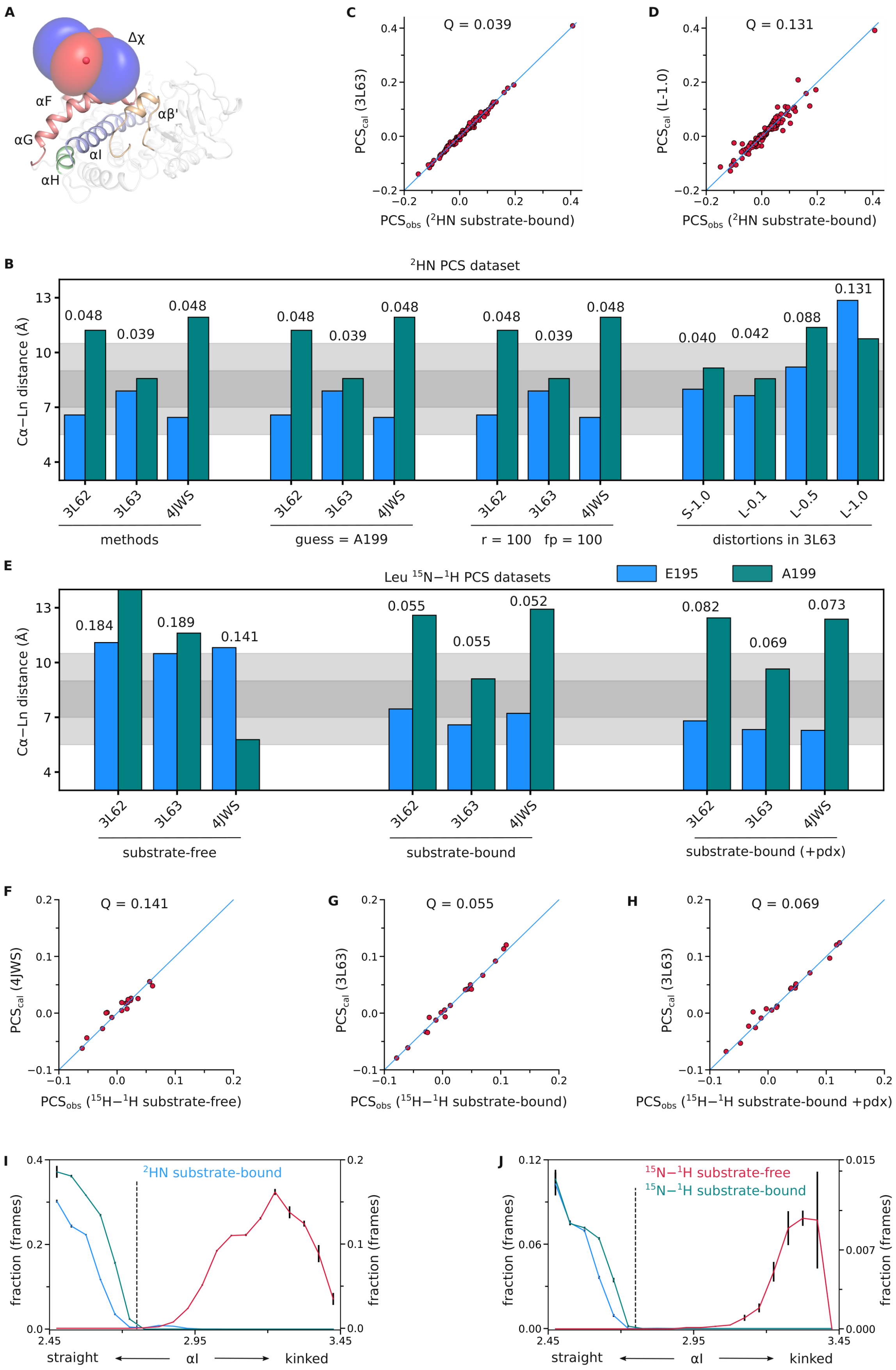
A
Δχ
αF
αG
αI
αH
αβ'
C
Q = 0.039
PCS_cal (3L63)
PCS_obs (²HN substrate-bound)
D
Q = 0.131
PCS_cal (L-1.0)
PCS_obs (²HN substrate-bound)
B
²HN PCS dataset
Cα−Ln distance (Å)
0.048
0.039
0.048
0.048
0.039
0.048
0.048
0.039
0.048
0.040
0.042
0.088
0.131
3L62
3L63
4JWS
methods
guess = A199
r = 100 fp = 100
S-1.0
L-0.1
L-0.5
L-1.0
distortions in 3L63
E
Leu ¹⁵N−¹H PCS datasets
E195
A199
0.184
0.189
0.141
0.055
0.055
0.052
0.082
0.069
0.073
substrate-free
substrate-bound
substrate-bound (+pdx)
F
Q = 0.141
PCS_cal (4JWS)
PCS_obs (¹⁵H−¹H substrate-free)
G
Q = 0.055
PCS_cal (3L63)
PCS_obs (¹⁵H−¹H substrate-bound)
H
Q = 0.069
PCS_cal (3L63)
PCS_obs (¹⁵H−¹H substrate-bound +pdx)
I
²HN substrate-bound
fraction (frames)
straight
αI
kinked
J
¹⁵N−¹H substrate-free
¹⁵N−¹H substrate-bound
fraction (frames)
straight
αI
kinked

**Figure S2:** (A) The estimated lanthanoid (Ln) ion and $\Delta\chi$ tensor, estimated by fitting $^{2}HN$ PCS with 3L63 crystal structure. (B) The $C_\alpha - Ln$ distances for residues E195 and A199, measured by various fitting schemes of $^{2}HN$ PCS data with crystal structures 3L62, 3L63, 4JWS and various distortions of 3L63. The value on top of bars are Q-scores. The S-1.0 to L-1.0 represents increasing distortions of 3L63**[1]. The dark and lightly shaded bands represents the 7-9 and 5.5-10.5 $\mathring{A}$ distances. (C,D) The matching between observed $^{2}HN$ PCS with that of calculated PCS from 3L63 and its L-1.0 distortion. (E) The $C_\alpha - Ln$ measured by fitting Leu $^{15}N -^{1}H$ PCS measurements with different crystals. (F-H) The matching observed and calculated PCS of Leu $^{15}N -^{1}H$ PCS with their correctly fitted crystal. (I,J) The fitting between MD structures with different PCS measurements, as done in maintext (Figure 3) but with cutoff 6-10 and 7-9 $\mathring{A}$ respectively. The vertical dashed line demarcates straight vs kinked conformations of $\alpha$I.

---

#### Supplementary note I: Benchmarking for NMR-PCS

We started with 4 NMR-PCS datasets as described in methods, i.e., (i) $^{2}HN$ PCS of substrate-bound P450cam, (ii-iv) Leu $^{15}N -^{1}H$ PCS of substrate-free, substrate-bound and substrate-pdx-bound P450cam respectively. We used crystal structures corresponding to pdb ids: 3L62 (open conformation), 3L63 (closed conformation) and 4JWS (open conformation) for benchmarking. As per observations by Ubbink and co-workers,[1] $^{2}HN$ PCS matches with 3L63 and not 3L62, $^{15}N -^{1}H$ substrate-free PCS matches with 3L62 and not 3L63, whereas $^{15}N -^{1}H$ substrate-bound and substrate-pdx-bound PCS matches with 3L63 and not 4JWS.

Pseudocontact shifts are measured as change in chemical shifts in presence of paramagnetic tag (Ln atom). This change can be quantified by relative positions of atoms-of-interest from paramagnetic tag and a magnetic susceptibility tensor $\Delta\chi$.[2] Given a set of PCS (observed in experiments) and atomic positions (crystal or MD structure), the $\Delta\chi$ tensor and position of Ln atom can be determined by iteratively minimizing the difference between observed and calculated PCS.[2] The paramagnetic tag is attached to two protein residues (E195 and A199 of $\alpha$G) via a CLaNP-7 linker. Therefore, for a correct fitting process, the fitted position of Ln atom should be nearby its attached residue, which was estimated to be 7-9 Å in previous works.[3] This $C\alpha - Ln$ distance is used as goodness of fit.

The $^2HN$ substrate-bound PCS were fitted with 3L62, 3L63 and 4JWS crystal structures, as per hyperparameter settings defined in methods. The estimated positions of Ln atom is within 7-9 Å for only 3L63 and not 3L62 or 4JWS (Figure S2A-C), in agreement with previous observation that $^2HN$ PCS are compatible with 3L63.[1] The change in hyperparameters $initia\_guess = A199C\alpha$ or $radius(r) = 100$, $fitting\_points(fp) = 100$ does not change the results, indicating the optimized hyperparameters (Figure S2B). If the atomic positions of 3L63 are distorted randomly**1, the goodness of fit starts decreasing for small distortions while do not fit ($C\alpha - Ln$ distance ¿ 7-9 Å) for large distortions (Figure S2B,D).

The Leu $^{15}N -^1 H$ substrate-free, substrate-bound, substrate-pdx-bound PCS were fitted with 3L62, 3L63 and 4JWS crystal structures. None of the crystals matched with any of the PCS measurements, if a strict cutoff of $C\alpha - Ln$ distance within 7-9 Å is considered (Figure S2E). Nevertheless, the expected matching of $^{15}N -^1 H$ substrate-free with 4JWS and both $^{15}N -^1 H$ substrate-bound with 3L63 can be attained if a tolerable cutoff of $C\alpha - Ln$ distance of 5.5-10.5 Å is considered (Figure S2E-H). The strict cutoff of 7-9 Å might not be very applicable to structures, given that crystals generally have atomic resolution of the order of 1-2 Å. Similarly, MD structures have fast thermal fluctuations of the order of 3-4 Å. It is in agreement, that previous acceptable PCS matching with structural ensembles did not strictly fall between 7-9 Å.[4,5] Hence considering the crystal resolutions and previous studies, the tolerable cutoff of 5.5-10.5 Å was used as goodness of fit. This tolerable cutoff attains the expected fitted patterns of PCS with crystals. Further, the results in maintext (Figure 3) does not does not depend on cutoff used, as the same trends are achievable with other cutoffs (Figure S2I,J).

****1: Distortions of crystal structure 3L63**: The distortions in the HN atomic positions of 117 residues corresponding to $^2HN$ PCS were created to remove their observed fitting (as a negative control). Small (S) and large (L) distortions were created by randomly changing the tenths or ones place respectively of anyone of the x, y or z coordinates. The small distortions were created for all atomic positions (S-1.0). Whereas the large distortions were created 10%, 50% or all atomic positions, represented as L-0.1, L-0.5 and L-1.0 respectively.

---

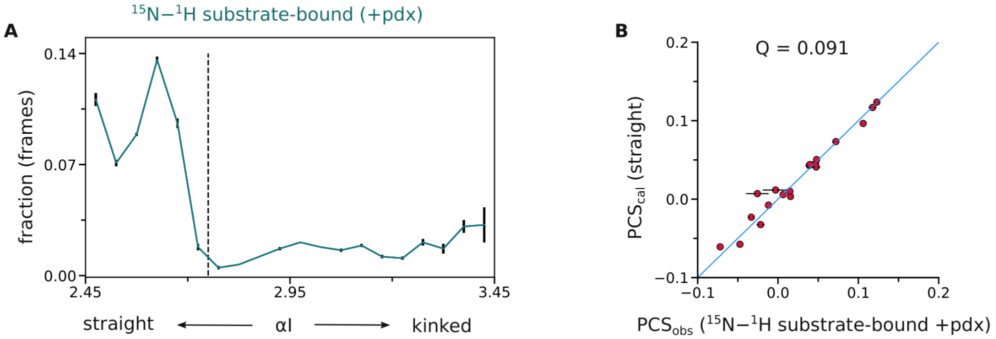


**Figure S3:** (A) The fraction of MD structures correctly fitted with Leu $^{15}-^{1}H$ PCS of substrate-bound (+pdx) state. The vertical dashed line demarcates straight vs kinked conformations of $\alpha$I. (B) The matching between observed and MD calculated PCS.

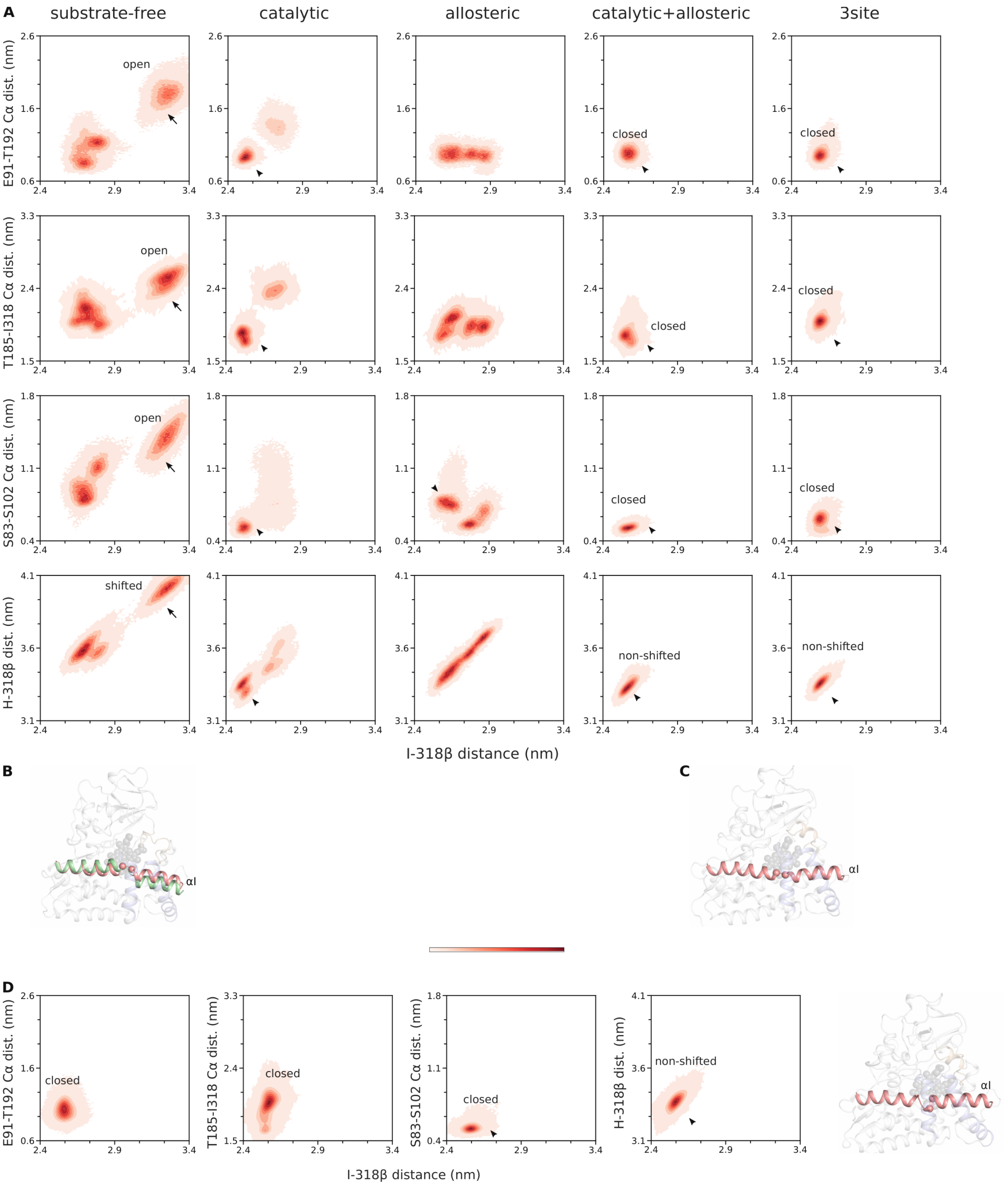


**Figure S4:** The conformational correlation plots of channel-1, channel-2 and $\alpha$H with $\alpha$I measured on previously reported simulations of different substrate-bound stages of P450cam+pdx complex[5] (A-C), substrate-binding simulations of closed P450cam[6] (D). The conformational correlation is in agreement with here-described allosteric mechanism, except for P450cam+pdx-allosteric state where intermediate kink in $\alpha$I is not followed with intermediate opening in channel-1 (remains closed). The snapshots in (B) and (C) are representative of P450cam+pdx-substrate-free and P450cam+pdx-catalytic+allosteric simulation ensembles respectively. The previous simulation data after 350 ns were used. The arrowhead and arrow represents the straight and kinked conformations of $\alpha$I.

---

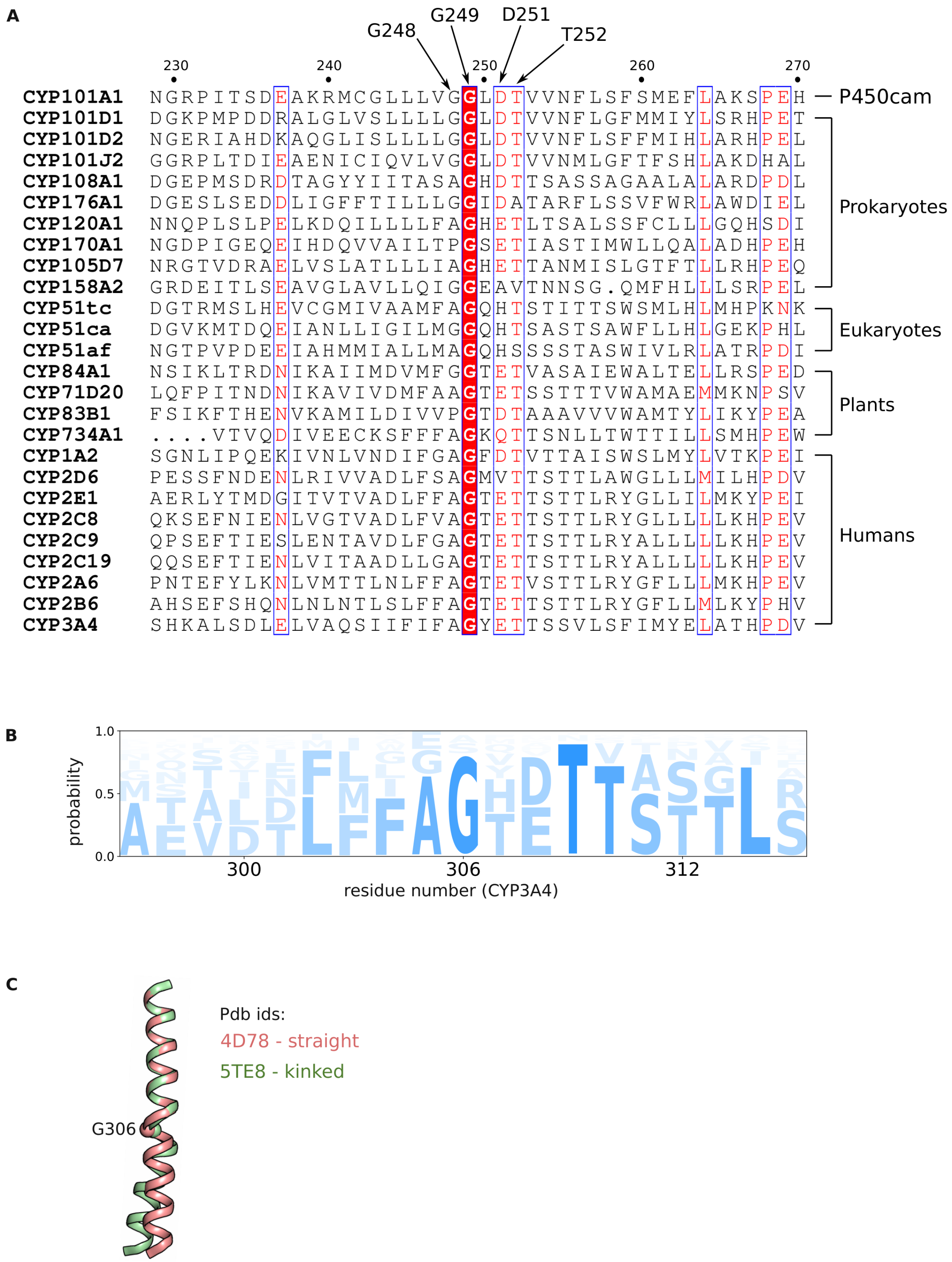
A
G248
G249
D251
T252
230
240
250
260
270
CYP101A1 NGRPITSDEAKRMCGLLLVGGLDTVVNFLSFSMEFLAKSPEH
CYP101D1 DGKPMPDDRALGLVSLLLLGGLDTVVNFLGFMMIYLSRHPET
CYP101D2 NGERIAHDKAQGLISLLLLGGLDTVVNFLSFFMIHLARHPEL
CYP101J2 GGRPLTDIEAENICIQVLVGGLDTVVNMLGFTFSHLAKDHAL
CYP108A1 DGEPMSDRDTAGYYIITASAGHDTTSASSAGAALALARDPDL
CYP176A1 DGESLSEDDLIGFFTILLLGGIDATARFLSSVFWRLAWDIEL
CYP120A1 NNQPLSLPELKDQILLLLFAGHETLTSALSSFCLLLGQHSDI
CYP170A1 NGDPIGEQEIHDQVVAILTPGSETIASTIMWLLQALADHPEH
CYP105D7 NRGTVDRAELVSLATLLLIAGHETTANMISLGTFTLLRHPEQ
CYP158A2 GRDEITLSEAVGLAVLLQIGGEAVTNNSG.QMFHLLLSRPEL
CYP51tc DGTRMSLHEVCGMIVAAMFAGQHTSTITTSWSMLHLMHPKNK
CYP51ca DGVKMTDQEIANLLIGILMGGQHTSASTSAWFLLHLGEKPHL
CYP51af NGTPVPDEEIAHMMIALLMAGQHSSSSTASWIVLRLATRPDI
CYP84A1 NSIKLTRDNIKAIIMDVMFGGTETVASAIEWALTELLRSPED
CYP71D20 LQFPITNDNIKAVIVDMFAAGTETSSTTTVWAMAEMMKNPSV
CYP83B1 FSIKFTHENVKAMILDIVVPGTDTAAAVVVWAMTYLIKYPEA
CYP734A1 ....VTVQDIVEECKSFFFAGKQTTSNLLTWTTILLSMHPEW
CYP1A2 SGNLIPQEKIVNLVNDIFGAGFDTVTTAISWSLMYLVTKPEI
CYP2D6 PESSFNDENLRIVVADLFSAGMVTTSTTLAWGLLLMILHPDV
CYP2E1 AERLYTMDGITVTVADLFFAGTETTSTTLRYGLLILMKYPEI
CYP2C8 QKSEFNIENLVGTVADLFVAGTETTSTTLRYGLLLLLKHPEV
CYP2C9 QPSEFTIESLENTAVDLFGAGTETTSTTLRYALLLLLKHPEV
CYP2C19 QQSEFTIENLVITAADLLGAGTETTSTTLRYALLLLLKHPEV
CYP2A6 PNTEFYLKNLVMTTLNLFFAGTETVSTTLRYGFLLLMKHPEV
CYP2B6 AHSEFSHQNLNLNTLSLFFAGTETTSTTLRYGFLLMLKYPHV
CYP3A4 SHKALSDLELVAQSIIFIFAGYETTSSVLSFIMYELATHPDV
P450cam
Prokaryotes
Eukaryotes
Plants
Humans
B
1.0
0.5
0.0
probability
300
306
312
residue number (CYP3A4)
C
Pdb ids:
4D78 - straight
5TE8 - kinked
G306

**Figure S5:** (A) A multiple sequence alignment of representative sequence of cytochrome P450 from different kingdoms, including the pharmaceutically most important P450s of humans. Residue numbering is according to CYP101A1. (B) The concensus sequence of $\alpha$I region for human cytochrome P450 sequences. The residue numbering is according to CYP3A4. (C) The representative crystal structures of human CYP3A4, highlighting straight and kinked conformations of $\alpha$I, indicating the potential existence of hereby-defined $\alpha$I conformations upto humans.

---

**Table S1:** The pdb ids classified into straight and kinked conformations of $\alpha$I. The straight conformations possessed channel-1 and channel-2 in closed conformation, while in open conformation with kinked conformation of $\alpha$I. The pdb ids corresponding to exceptions due to linkered substrates are also shown. Note: Pdb id 1GJM is exception in straight conformation, as it contains beta sheet in place of $\beta$' helix (part of channel-2). The pdbs of straight conformation are almost same with each other except few differences in flexible loops between $\alpha$G-$\alpha$H and $\alpha$H-$\alpha$I.

| Straight conformation | Kinked conformation |
|---|---|
| 1AKD, 1C8J, 1CP4, 1DZ4, 1DZ6, 1DZ8, 1DZ9, 1GJM 1IWI, 1IWJ, 1IWK, 1J51, 1MPW, 1NOO, 1O76, 1P2Y, 1P7R, 1PHA, 1PHB, 1PHC, 1PHD, 1PHE, 1PHF, 1PHG, 1T85, 1T86, 1T87, 1T88, 1UYU, 1YRD, 2A1M, 2A1N, 2CP4, 2CPP, 2FE6, 2FER, 2FEU, 2FRZ, 2GQX, 2GR6, 2H7Q, 2H7R, 2H7S, 2L8M, 2LQD, 2M56, 2QBL, 2QBM, 2QBN, 2QBO, 2Z97, 2ZAW, 2ZUH, 2ZUI, 2ZUJ, 2ZWT, 2ZWU, 3CP4, 3CPP, 3FWF, 3FWG, 3FWI, 3FWJ, 3L63, 3WRH, 3WRJ, 3WRL, 3WRM, 4CP4, 4CPP, 4EK1, 4G3R, 4KKY, 4L49, 4L4A, 4L4B, 4L4C, 4L4D, 4L4E, 4L4F, 4L4G, 5CP4, 5CPP, 5WK7, 5WK9, 6CP4, 6CPP, 6WE6, 6WFL, 7CPP, 8CPP | 1K20, 1QMQ, 3L61, 3L62, 3P6Q, 3P6R, 3P6S, 3P6U, 3P6V, 3P6W, 3P6X, 3W9C, 3WRI, 3WRK, 4JWS, 4JWU, 4JX1, 5GXG, 5IK1, 6NBL<br><br>Exceptions by linkered substrates<br>1LWL, 1RE9, 1RF9, 3OIA, 3OL5, 3P6M, 3P6N, 3P6O, 3P6P, 3P6T |

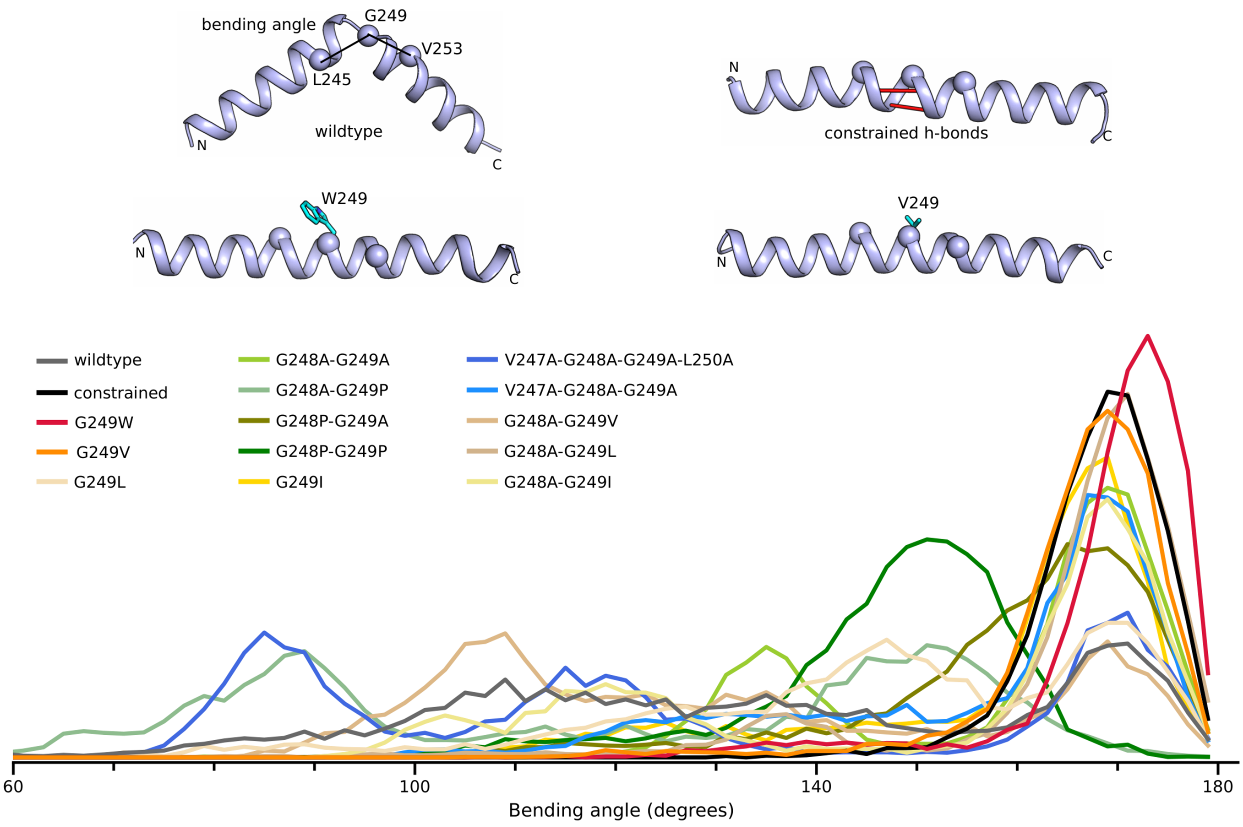


**Figure S6:** Top panel exhibits snapshots from wildtype, constrained h-bonds, G249W and G249V variants of simulations, highlighting the bending angle calculation (black lines in wildtype) and constrained h-bonds (red lines). Bottom plot represents the probability distribution of bending angle as observed for different variants of $\alpha$I.

#### Supplementary note II: Searching for I helix mutants

With the objective of finding a $\alpha$I mutant with stable straight conformation, the unbiased MD simulations of trail system, constituting only $\alpha$I (residues 234-266), were performed (methods). Compared to complete P450cam simulation ensembles, the only $\alpha$I simulations demonstrate a very large bending, as it is now free of 3D-structural constraints. This bending centered at G249 which is also the source of straight-kinked conformations (Figure S6, top panel), and can be used as proxy for kinked conformation. This bending can be quantified by angle defined by $C\alpha$ atoms of residues L245, G249 and V253. In simulations, the wildtype $\alpha$I samples all possible bend-to-straight conformations (Figure S6, bottom). As an alternative, a constrained system was designed to sample only straight conformation. The broken backbone hydrogen (h-)bonds by G248-G249 residues were re-enforced by putting distance contraints between L246-L250 and V247-D251 residue pairs (Figure S6, top). Though un-physical, the constrained system was able to sample straight conformations with bending angle close to $180^o$ (Figure S6). Using the wildtype and contrained systems as negative and positive controls respectively, different $\alpha$I mutants were simulated, with the objective to match the bending angle positive control. The high helix propensity alanine substitutions, very rigid proline substitutions, their combinations, to other hydrophobic residue substitutions were attempted. The observations of this unphysical trial system were transferable to full P450cam system. For instance, the only $\alpha$I with G248A-G249A mutations sample majorly straight and minorly bend conformations, which were equivalent to majorly straight and minorly kinked conforamtions observed for full G248A-G249A-P450cam (maintext). Nevertheless, the trial system was only used for leads and ultimately checked for full P450cam system. The bending angles of different only $\alpha$I mutants indicate that G249W sampled predomiant straight conformations, close to $180^o$ and even better than positive control system (Figure S6). G249V and G248A-G249L also sampled straight conformations as good as positive control, out of which G249V was selected, owing to single mutation.

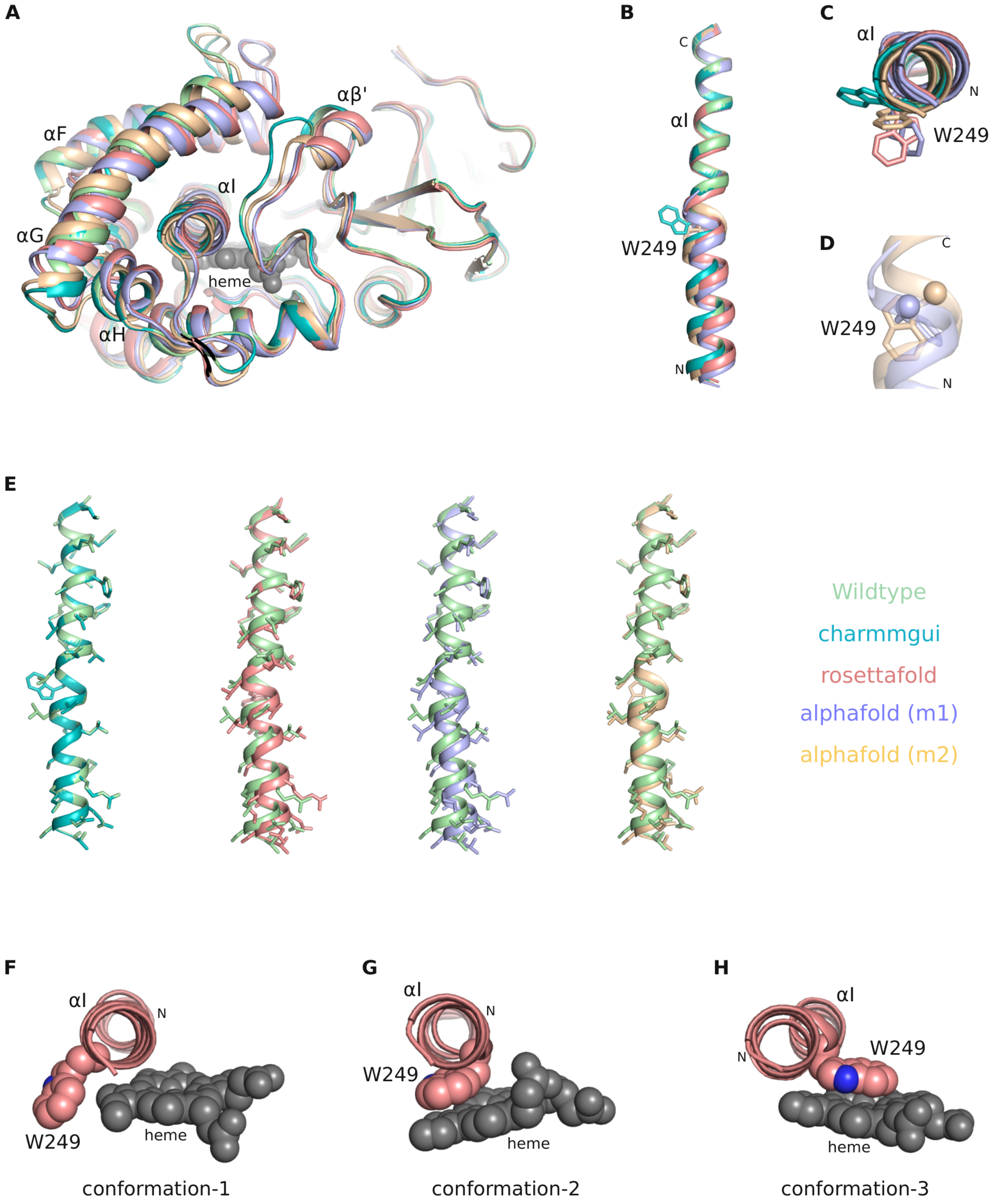


**Figure S7:** (A-E) The starting structures of G249W-substrate-free P450cam generated by different algorithms, aligned with wildtype structure, and zoomed views of $\alpha$I. Compared to other structures, the residue W249 is shifted towards heme side in second best model of alphafold (m2, D snapshot), which results in different samplings in simulations. (F-H) Different conformations of W249 observed in simulations. W249 remain sideways in G249-1 simulation ensemble, without any steric interference with heme moiety, as in (F) snapshot. The simulation ensemble started from alphafold-second best model (m2), which have W249 shifted towards heme side (D snapshot) samples two conformations as shown in (G) and (H) snapshots. In (G) conformation with major population, the W249 underlies the $\alpha$I in tolerable conformation. While in minor (H) conformation, the W249 covers the heme moiety and $\alpha$I in kinked conformation. The (H) conformation might not be a feasible conformation, but it might be the consequence of choosing second best model of alphafold. The (F) conformation has no impact on heme.

---

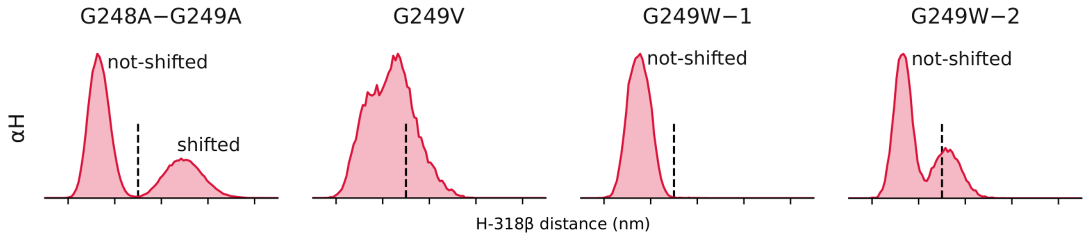


**Figure S8:** The $\alpha$H conformation in different mutants of $\alpha$I. The not-shifted conformations of $\alpha$H were observed to the extent straight conformation of $\alpha$I was achieved by the mutant (maintext).

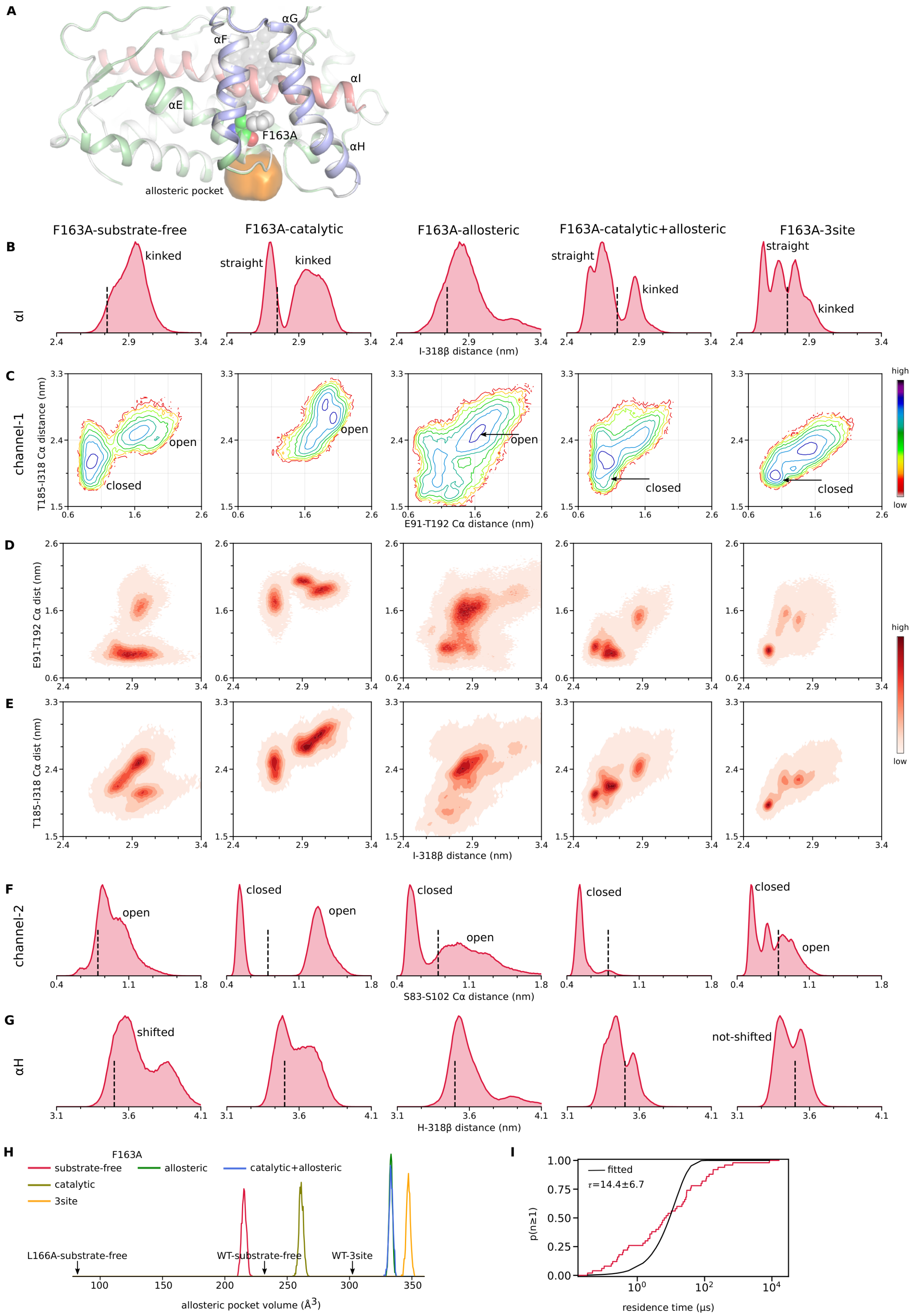
A
αG
αF
αI
αE
F163A
αH
allosteric pocket
B
F163A-substrate-free
F163A-catalytic
F163A-allosteric
F163A-catalytic+allosteric
F163A-3site
αI
kinked
straight
I-318β distance (nm)
C
channel-1
T185-I318 Cα distance (nm)
open
closed
E91-T192 Cα distance (nm)
high
low
D
E91-T192 Cα dist (nm)
E
T185-I318 Cα dist (nm)
I-318β distance (nm)
F
channel-2
S83-S102 Cα distance (nm)
G
αH
shifted
not-shifted
H-318β distance (nm)
H
F163A
substrate-free
allosteric
catalytic+allosteric
catalytic
3site
L166A-substrate-free
WT-substrate-free
WT-3site
allosteric pocket volume (Å$^3$)
I
fitted
τ=14.4±6.7
p(n≥1)
residence time (μs)

**Figure S9:** (A) The starting structure of F163A mutant P450cam overlayed on wildtype starting structure (white cartoon). The F163A mutation (white to green spheres) and allosteric pocket (orange surface) are highlighted. (B,C) The $\alpha$I and channel-1 conformations in different substrate-bound stages of F163A P450cam. (D,E) The conformation correlation between $\alpha$I and channel-1 conformations in different substrate-bound stages. (F,G) The channel-2 and $\alpha$H conformations in different substrate-bound states. (H) The allosteric pocket volume different substrate-bound stages of F163A P450cam. The arrow indicates allosteric pocket volumes in wildtype and L166A P450cam. (I) The ECDF of metadynamics drived unbinding times and fitted $P(n \geq 1)$ curve for F163A-allosteric mutant. The y-axis in (B,F,G,H) and red-to-blue in (C) and white-to-red in (D,E) represent arbitrary probability axis.

---

#### Supplementary note III: Decoupling channel-1 from I helix

We attempted to design a decoupled P450cam as a negative proof of allosteric mechanism, where $\alpha$I does not possessed conformational correlation with other structural elements. We emphasized on channel-1, such that in decoupled P450cam, its closed-open conformations would not be correlated with straight-kinked conformations of $\alpha$I. We surmised that allowing enough space for $\alpha$I to attain kinked conformation comfortably within structural fold of P450cam, might decoule it from other structural elements. For instance, creating enough space towards $\alpha$FG side of $\alpha$I might decouple it from channel-1. Towards this end, F163 of $\alpha$E and between $\alpha$FG and $\alpha$I was mutated to small alanine residue (Figure S9A).

In F163A-substrate-free state, the $\alpha$I expectedly sampled predominant kinked conformation owing to extra space allowed by F163A mutation (Figure S9B). As per allosteric mechanism, the channel-1 should sample predominant open conformation. Conversely, the channel-1 in F163A-substrate-free state sampled both open and closed conformations with minima in closed conformation (Figure S9C). The conformational correlation plots exhibit no correlation between $\alpha$I and channel-1 in F163A-substrate-free state (Figure S9D,E). The channel-2 and $\alpha$H sampled predominant open and shifted conformations respectively in F163A-substrate-free state, inline with allosteric mechanism (Figure S9F,G). Overall in substrate-free state, the F163A mutant successfully possesed channel-1 decoupled from $\alpha$I, while channel-2 and $\alpha$H maintains their conformational correlation with $\alpha$I.

The successive addition of three substrate binding modes were able to shift $\alpha$I towards straight conformation (Figure S9B), though not completely as observed for WT-3site, but removing its decoupling with channel-1. In F163A-catalytic state, $\alpha$I sampled minorly straight and majorly kinked conformation (Figure S9B), inline with allosteric mechanism whereby catalytic mode hydrobhopically stabilize straight conformation of $\alpha$I (Figure 2). As per allosteric mechanism, the channel-1 should sample minorly closed and majorly open conformation, instead F163A-catalytic state sampled channel-1 in predominant open conformation only (Figure S9C). The conformational correlation plots indicate that decoupling between channel-1 and $\alpha$I largely persists in F163A-catalytic (Figure S9 D,E). In F163A-allosteric state, the $\alpha$I slightly shifts towards straight conformation (compared to F163A-substrate-free) but largely maintained its kinked conformation (Figure S9B). The channel-1 sampled mainly open conformation but also closed conformation (Figure S9C) indicating that decoupling persists, but conformational correlation between channel-1 and $\alpha$I starts to appear in F163A-allosteric state (Figure S9D,E). In presence of allosteric and other active site binding modes (F163A-catalytic+allosteric, F163A-3site), the channel-1 sampled majorly closed and minorly open conformations in accordance with majorly straight and minorly kinked conformations of $\alpha$I (Figure S9B,C). The conformational correlation plots indicate significant correlation between channel-1 and $\alpha$I as observed for wildtype (Figure S9D,E).

The decoupling between channel-1 and $\alpha$I of F163A-substrate-free state is lost upon addition of substrates, specifically with the allosteric mode. Compared to F163A mutation site , the allosteric site lies on the opposite side of $\alpha$E and some of the empty space created by mutation leaked to allosteric site. For instance, despite the predominant shifted conformation of $\alpha$H, the allosteric pocket in F163A-substrate-free maintained its volume of 215.3±1.9 $\AA^3$ which is close to WT-substrate-free (234.8±1.8 $\AA^3$) and much larger than L166A-substrate-free (82.4±1.1 $\AA^3$) (Figure S9H). Similarly in substrates bound F163A-3site, the allosteric site is very large with pocket volume of 347.1±1.6 $\AA^3$, even larger than WT-3site (303.9±2.1 $\AA^3$). To re-confirm, the residence time of allosteric mode in F163A-allosteric with predominantly shifted $\alpha$H was measured, which comes out to be 14.4±6.7 $\mu$s (Figure S9I), more than 4.2±0.2 $\mu$s in wildtype.[5] Therefore, we concluded that the empty space created by F163A mutation atleast partially leaks to allosteric site, which is taken up by allosteric mode and hence removed the decoupling. Overall, F163A is decoupled P450cam only in substrate-free state and not in substrate-bound states. The channel-2 and $\alpha$H maintained their coupling with $\alpha$I (Figure S9E,G).

---

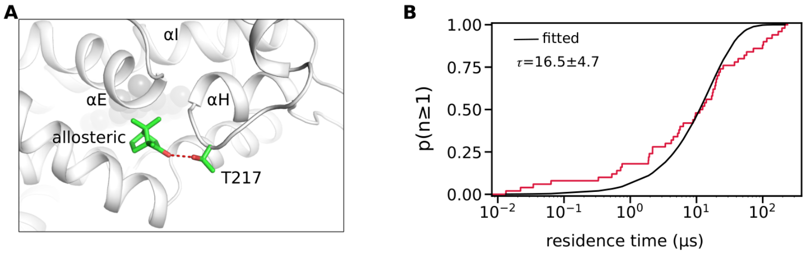


**Figure S10:** (A) Zoomed view of allosteric mode hydrogen bonded to T217. (B) The ECDF of metadynamics drived unbinding times and fitted $P(n \geq 1)$ curve for T217V-allosteric mutant.
